# Cohesin mutation sensitizes cancer cells to anti-PD-1 therapy through endogenous retrovirus-mediated PD-L1 upregulation

**DOI:** 10.1101/2022.02.19.481125

**Authors:** Yumin Han, Fangfei Peng, Yunqi Chang, Tingting Liu, Jiayan Shen, Zizhuo Chen, Qian Dong, Ping Zhou, Feng Jiang, Honggang Xiang, Hong Zhu, Chen Qing, Xiangyin Kong, Jian Ding, Jing-Yu Lang

## Abstract

Immune checkpoint therapy shows impressive and durable clinical responses in cancer patients, but the genetic determinants that enable cancer cells to respond to anti-PD-1 therapy are still elusive. Herein, we identified that *NIPBL* deficiency promotes endogenous retrovirus (ERV) expression in tumour cells, which in turn inactivates CD8+ tumour-infiltrating lymphocytes (TILs) via the PD-L1/PD-1 inhibitory checkpoint pathway. Mechanistically, *NIPBL* deficiency impairs DNMT1 transcription, preventing DNMT1 from suppressing ERV expression in tumour cells; ERVs stimulate PD-L1 expression by inducing the STAT2-IRF9 complex, a downstream event of double-stranded RNA (dsRNA)-MAVS-IRF3 signalling, and thereby suppress CD8 TIL-mediated immunity. An anti-PD-1 monoclonal antibody achieved remarkable therapeutic effects in *Nipbl*-deficient syngeneic tumour models and improved host survival by eliciting an antitumour memory immune response. Cancer patients harbouring mutations of cohesin subunits and regulators plus DNMT1 had significantly better responses to anti-PD-1 therapy than their non-mutated counterparts did. Our study reveals a novel mechanism by which cohesin complex deregulation stimulates ERV expression by impairing DNMT1 expression and fosters an immunosuppressive tumour microenvironment by activating the PD-L1/PD-1 inhibitory checkpoint.

## Main

Immune inhibitory checkpoint molecules play a critical role in tumour development. For example, programmed cell death ligand-1 (PD-L1) is a ligand for the PD-1 receptor, which is highly expressed in tumours. PD-L1 blocks the activation of T cell receptors when it binds to the PD-1 receptor and eventually enables tumour cells to escape from the antitumour immune response mediated by cytotoxic CD8+ T cells^1–3^. Anti-PD-1 monoclonal antibodies (mAbs), such as nivolumab and pembrolizumab, have achieved remarkable, persistent clinical outcomes in patients with melanoma, non-small-cell lung cancer and tumours with microsatellite instability (MSI)^4–7^. However, only a limited subpopulation of patients benefit from immune checkpoint block therapies^8, 9^. The genomic determinants of the response to anti-PD-1 therapy remain unknown.

To explore the genetic determinants for responding to anti-PD-1 therapy, we performed an *in vivo* genome-scale CRISPR-Cas9 knockout (GeCKO) screening, which revealed that loss of cohesin subunits and regulators increased tumour sensitivity to anti-PD-1 therapy by stimulating PD-L1 expression in tumour cells. Cohesin is a multiple subunit complex composed of core subunits (SMC1A/SMC1B, SMC3, STAG1/STAG2 and RAD21) and regulators (NIPBL, PDS5A/PDS5B, WAPL, CDCA5 and MAU2)^10^. NIPBL is a key factor for loading the cohesin complex at promoters. Apart from its role in sister chromatid cohesion, the cohesin complex has been implicated in 3D genome organization and gene regulation^11–15^. Recently, pan-cancer atlas analysis identified the cohesin complex as one of 16 most commonly mutated subnetworks in human cancer^15, 16^. However, it is unclear whether the cohesin complex functions by regulating immunity in cancer, and the immune-regulatory role of the cohesin complex remains largely unknown.

In this study, we reveal that knockout of *Nipbl* and other cohesin complex members stimulates dsRNA ERV expression by blocking DNMT1 transcription, which in turn boosts PD-L1 expression by activating the dsRNA-MAVS-IRF3-STAT2/IRF9 signalling pathway in tumour cells. Blockade of PD-1 using a specific monoclonal antibody resulted in 70% of *Nipbl*-deficient tumours completely regressing, remarkably improving mouse survival by reactivating cytotoxic CD8+ T cells. Mutations in cohesin subunits and regulators plus DNMT1 can predict the survival rate of melanoma, lung and colorectal cancer patients compared with that of their non-mutated counterparts. In summary, our study reveals a novel mechanism by which loss of cohesin subunits and regulators in tumours shapes the tumour-immune cell interaction and indicates that tumours with these mutations could be selectively treated with anti-PD1 therapy.

## Results

### *In vivo* genome-scale screening reveals that *Nipbl*-deficient tumours potentially respond to anti-PD-1 therapy

To identify genetic determinants in the response to anti-PD-1 therapy, we selected an immunotherapy-insensitive murine CT26 colorectal cancer cell line to perform GeCKO screening *in vivo*^17–19^. CT26 cells were infected with lentivirus containing a library of single guide RNAs (sgRNAs) at a multiplicity of infection (MOI) of 0.3 and then selected the cells for 7 days with puromycin to remove uninfected cells. The transduced cells were subcutaneously injected into the flanks of syngeneic mice (2×10^7^ cells per injection, 10 mice per group), which were then intraperitoneally administered either isotype IgG or anti-PD-1 monoclonal antibody five times (200 μg/mouse per injection, Fig. 1A). After treatment stopped, genomic DNA was extracted from each tumour sample, normalized and subjected to barcoded deep sequencing (see more information in the Material and Methods).

**Figure 1.**
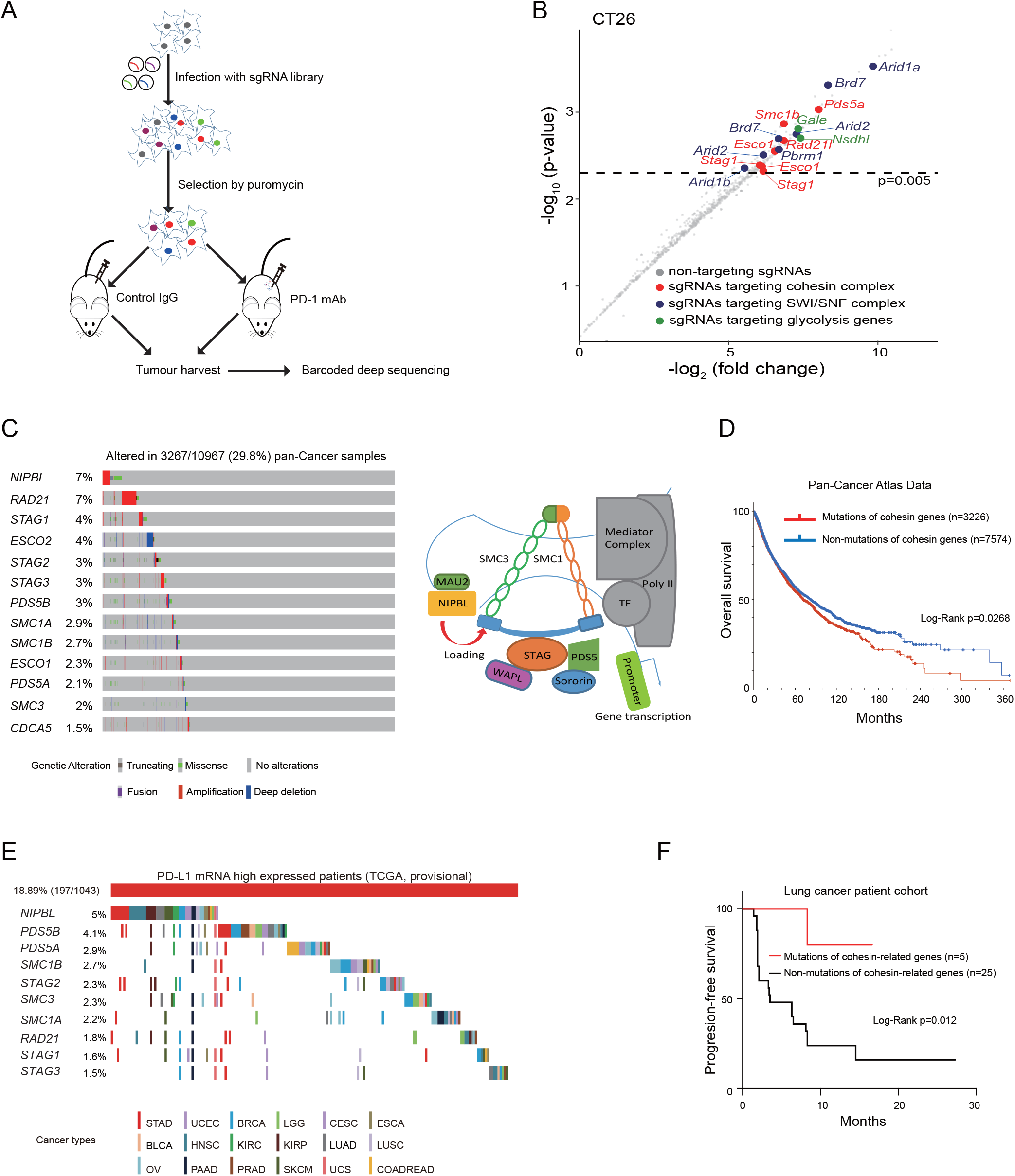
*In vivo* GeCKO screening reveals that loss of cohesin subunits and regulators sensitizes tumour cells to anti-PD-1 therapy. (A) Schematic diagram of the *in vivo* GeCKO screening strategy. (B) Compared to non-targeting sgRNAs, sgRNAs targeting cohesin subunits and regulators were significantly reduced by comparing the anti-PD-1 group to control IgG group. The dashed lines indicate a p value of 0.005. Annotated genes represent cohesin subunits and regulators (red), SWI/SNF complex (blue) and glycolysis (green) genes validated by other groups, and genes with p values > 0.005 (grey). (C) The mutation frequencies of 13 cohesin subunits and regulators in the pan-cancer atlas samples (n=10967). Cartoon of cohesin complex subunits and regulators. (D) Mutations of 13 cohesin subunits and regulators negatively correlated with the patient’s overall survival of pan-cancer samples. (E) The distribution of inactive mutations and null expression (log_2_ (RPKM) < -2) of the indicated cohesin subunits and regulators in PD-L1-positive (log2 (RPKM) > 0.5) tumour samples of patients across 18 cancer types from the TCGA database. (F) Mutations in *NIPBL*, *STAG2*, *STAG3* and *SMC1B* significantly prolonged the progression-free survival of lung cancer patients treated with pembrolizumab (n=5) compared with patients with non-cohesin complex member mutations (n=25) (log-rank p=0.012). The survival significance was assessed by the log-rank Mantel-Cox test.

T cell-based CRISPR-Cas9 screens have been described by several laboratories. For example, mutations of the phosphatase *Ptpn2* and the G protein-coupled receptor *APLNR* confers tumour cells resistant to T-cell-mediated cytotoxicity^20, 21^; mutations of SWI/SNF complex genes sensitize tumour cells to immunotherapy^22^. Recently, Fli1 is reported as a key negative regulator of cytotoxic CD8+ T cells^23^. Our approach emphasized sensitive selection of sgRNAs depleted by an *in vivo* anti-PD-1 therapy, which allowed us to specifically identify cancer mutations conferring sensitivity to anti-PD-1 therapy.

After normalized by medium normalization, the significant distribution of raw read counts in different groups were calculated by the MAGeCK algorithm (false discovery rate <0.05)^24^, and the fold changes were calculated by comparing read counts of anti-PD-1 group to that of control IgG group (Fig. 1B). We used the following criteria to select sensitive candidates: 1) was one of the top 6,000 reduced sgRNAs by comparing the anti-PD-1 group to the control group; 2) was one of sixteen commonly mutated subnetworks in human cancer^16^; and 3) had a minimum of >3 members per complex hit in a single screen with p value < 0.005 and fold change <0.03. As a positive control, sgRNAs targeting the SWI/SNF complex (*Pbrm1*, *Brd7*, *Arid2, Arid1a* and *Arid1b*), which were previously reported to mediate the immune response to anti-PD-1 therapy, were significantly reduced in the anti-PD-1 group compared to the vehicle group (Figs. 1B and S1A) ^22^. As previously reported^20, 22^, glycolysis genes such as *Gale* and *Nsdhl* as well as the phosphatase *Ptpn2* were also obviously depleted in our screening (Fig. S1B and S1C). Strikingly, sgRNAs targeting cohesin subunits and regulators were significantly reduced in the anti-PD-1 group compared to that of the control IgG group (Fig. 1B). For example, sgRNAs targeting cohesin complex subunits (*Stag1, Smc1b* and *Rad21l)* and regulators (*Pds5a* and *Esco1*) were significantly reduced in the anti-PD-1/vehicle group compared to nontargeting sgRNA group (p<0.005, Fig. S1D). *Smc1b* and *Rad21l1* are meiotic-specific cohesin subunits, which are actively expressed in many tumour types^25, 26^.

As one of most commonly mutated cancer subnetworks, cohesin complex subunits and regulators were widely mutated in approximately 29.8% of 10,967 pan-cancer cases of thirty-two human cancer types and well correlated with patients’ overall, progression-free and disease-free survivals (Figs. 1C, 1D and S2)^15, 16, 27–29^. Notably, *NIPBL*, a key factor for loading the cohesin complex^11^, ranked as the top candidate with an approximately 7% mutation rate in all human cancer types. For *NIPBL*, there were 124 cases of amplification (copy number ≥ 4, 16.7%), 159 cases of truncation (21.4%), 424 cases of missense (57.1%) and other mutations. Remarkably, cohesin complex subunits and regulators were inactively mutated in tumour samples that were PD-L1-positive (log_2_ (RPKM) > 0.5) across eighteen cancer types from The Cancer Genome Atlas (TCGA), especially in stomach and colorectal tumours (Fig. 1E) ^27, 30^.

Patient survival analysis revealed that anti-PD-1 therapy significantly prolonged the survival rate of lung cancer patients with mutations of cohesin subunits and regulators compared to that of patients without mutations (p=0.012, Fig. 1F and Supplemental Table 1)^31^. These data indicate that deficiency of cohesin subunits and regulators such as *NIPBL* potentially sensitizes tumour cells to anti-PD-1 therapy.

### *Nipbl* loss facilitates tumour progression by inactivating CD8+ T cells through PD-1/PD-L1 inhibitory checkpoint

The inactivating mutation of *NIPBL* is well-documented with poorly characterization of immunomodulatory function^32–34^, so we chose it for further studies. Flow cytometry data illustrated that cell surface PD-L1 expression was significantly increased in CT26 (ratio=2.69, p=4.5E-5) and colon26 (ratio=4.02, p=1.1E-3) murine syngeneic colorectal tumour models when *Nipbl* was knocked out (Fig. 2A). Non-targeting sgRNA (sgControl) was used as a negative control. The PD-L1 mRNA and protein expression levels of sgNipbl CT26 and colon26 murine colorectal tumour cells were robustly increased compared to those of sgControl cells (Fig. 2B and 2C). Similar results were obtained using the HCT-116 human colorectal cancer cell line (Fig. 2D). The sgRNA-mediated knockout of Nipbl in CT26, colon26 and HCT-116 cells was validated by Sanger sequencing (Fig. S3A-S3C). Notably, high PD-L1-expressing B16F10 and cloudman S91 murine tumour cell lines had little or no detectable Nipbl protein expression compared to that of *Nipbl* wild-type CT26 cells (Fig. 2E) ^18^. *NIPBL^p.K603fs^*-mutant RKO human colorectal tumour cells also had more potent PD-L1 expression than *NIPBL* wild-type HCT-116 cells (Fig. 2F and see more cell lines in Fig. 5B).

**Figure 2.**
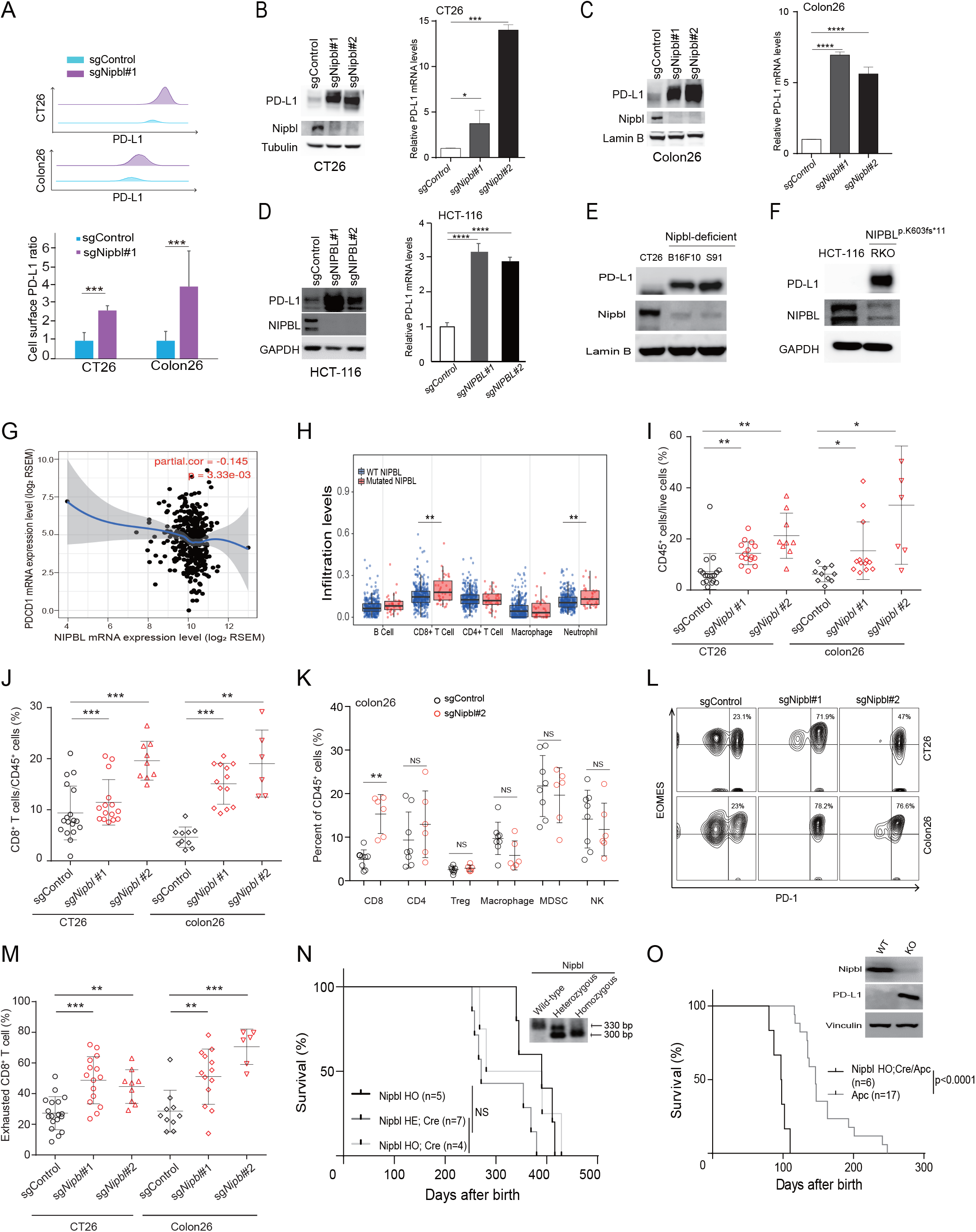
Nipbl loss suppresses tumour infiltrated CD8+ T cells by inducing the PD-L1/PD-1 inhibitory checkpoint pathway. (A) Flow cytometric analysis of PD-L1 protein expression in Nipbl knockout CT26 and colon26 syngeneic tumour models compared to sgControl models. The statistical bar diagram is shown on the bottom. (B-D) The indicated protein expression levels were detected by immunoblotting of CT26, colon26 and HCT-116 tumour cell lines, while the mRNA expression levels were determined by qRT-PCR. (E) PD-L1 and Nipbl protein expression was detected by immunoblotting in murine B16F10 and cloudman S91 tumour cell lines. (F) PD-L1 and NIPBL protein expression was detected in the human NIPBL^p.K603fs*11^ RKO colorectal tumour cell line. (G) Correlation of NIPBL mRNA levels in tumour cells with the PD-1 mRNA levels in infiltrating immune cells of TCGA colorectal cancer patient samples (n=457). Spearman’s correlation and estimated significance were calculated with the Tumor Immune Estimation Resource (TIMER) and adjusted for tumour purity. (H) Analysis of immune cell infiltration in NIPBL wild-type and NIPBL-mutant TCGA colorectal cancer samples (n=457). (I-J) The percentages of CD45+ cells and CD8+ TILs were quantitated (CT26 tumours: sgControl, n=18, sgNipbl#1, n=15, sgNipbl#2, n=9; colon26 tumours: sgControl, n=10, sgNipbl#1, n=13, sgNipbl#2, n=6). (K) The percentages of CD8+ and CD4+ TILs, Treg cells, macrophage, myeloid-derived suppressor cells (MDSCs) and natural killer (NK) cells were quantitated, normalized by CD45+ cells. (L) Representative plots showing the gating for the exhausted CD8+ T cell subpopulation characterized by PD-1+ and EOMES+ staining. (M) The percentages of exhausted CD8+ T cells in the indicated groups were quantitated (CT26 tumours: sgControl, n=18, sgNipbl#1, n=15, sgNipbl#2, n=9; colon26 tumours: sgControl, n=10, sgNipbl#1, n=13, sgNipbl#2, n=6). (N) The survival rates of Nipbl^fl/fl^ mice (n=5), Villin-Cre^+^; Nipbl^+/fl^ mice (n=7) and Villin-Cre^+^; Nipbl^fl/fl^ mice (n=4). For the loxP site in intron 4 of the Nipbl^fl/fl^ allele, the PCR products were 300 bp for the floxed allele and 330 bp for the wild-type allele. (O) The survival rates of APC^min^ mice (n=17) and Villin-Cre^+^; Nipbl^fl/fl^ mice; APC^min^ (n=6). The expression levels of the indicated proteins were determined by immunoblotting. The survival significance was assessed by the log-rank Mantel-Cox test. Data are shown as the mean ± SD, and the p value was calculated by two-tailed t tests.*p < 0.05, **p < 0.01, ***p < 0.001, ****p < 0.0001.

PD-L1 expression in tumour cells often lead to the exhaustion of CD8 TILs by activating the PD-1/PD-L1 inhibitory checkpoint^35^. Tumour-infiltrating immune cell data analysis^36^ revealed that NIPBL mRNA expression levels in tumour cells were negatively correlated with PD-1 mRNA levels in the infiltrating immune cells of TCGA colon adenocarcinomas (n=457, p=3.33E-3, Fig. 2G). Immune cell subtype analysis further revealed that the proportion of infiltrated CD8+ T cells was significantly increased in *NIPBL*-mutant colon tumour samples compared to *NIPBL* wild-type samples (p<0.01, Fig. 2H). These data suggest that *NIPBL* loss potentially inactivates CD8+ T cells by promoting the PD-1/PD-L1 interaction.

To verify this hypothesis, we inoculated *Nipbl* knockout murine tumour cells into the flanks of syngeneic mice and observed that the proportions of CD45+ and CD8 TILs were significantly increased in both sgNipbl CT26 and colon26 tumour tissues compared to sgControl tumour tissues (p<0.05, Fig. 2I and 2J); importantly, the portion of CD8 TILs but no other immune cell subtypes was significantly increased in sgNipbl tumour tissues compared to sgControl tumour tissues (p<0.01, Fig. 2K). Moreover, CD8 TILs expressing PD-1 but not Eomes, two markers of exhausted T cells^37^, were increased by two-to-threefold when *Nipbl* was knocked out (Figs. 2L, 2M and S4A), supporting the notion that *Nipbl* mutation blocks the antitumour immunity of CD8 TILs via the PD-1/PD-L1 inhibitory checkpoint pathway.

Noteworthy, we observed weakly or no expression of Cas9 and the phosphorylation of histone H2AX at Ser13 (γH2AX)^38^ in sgNipbl and sgControl CT26, colon26 and HCT-116 tumour cells (Fig. S5A and S5B). Exogenously overexpressed Cas9 and mitomycin C-induced γH2AX were used as positive controls. There were no significant change of the cytosolic dsDNA content of sgNipbl CT26 and colon26 compared to that of sgControl cells (p>0.05, Fig. S5C). Moreover, shSTING had no effect on PD-L1 expression compared to scramble (Fig. S5D). These data suggest that double-strand DNA breaks is not responsible for Nipbl loss-induced PD-L1 expression.

Next, we asked whether *Nipbl* loss promotes tumour growth in mice. Although cohesin knockout mice are early embryonic lethal^39, 40^, tissue-specific *Rad21* knockout mice has a normal lifespan^41^. Thus, we established *Villin*-Cre^+^; *Nipbl^flox/flox^* mice to achieve *Nipbl* deletion in gastrointestinal epithelial cells in which *NIPBL* is highly mutated^32, 42^. We observed no tumour formation in mice when heterozygous or homozygous *Nipbl* deletion was achieved in gastrointestinal epithelial cells; these mice showed a normal lifespan and pregnancy similar to that in control *Nipbl^flox/flox^* mice (p>0.05, Fig. 2N). When crossed with Apc^Min^ mice, *Nipbl* loss significantly promoted tumour progression by reducing the survival rate of these mice compared to that of Apc^Min^ mice, and this change was accompanied by increased PD-L1 expression (p<0.0001, Fig. 2O).

Collectively, we conclude that *NIPBL* loss promotes tumour progression by promoting the PD-L1/PD-1 interaction.

### *Nipbl* loss sensitizes tumour cells to anti-PD-1 therapy *in vivo* independent of tumour neoantigen

Herein, we examined whether *Nipbl*-deficient tumours were sensitive to anti-PD-1 therapy. As shown in Fig. 3A, *Nipbl*-deficient colon26 tumours were almost completely regressed 40 days after anti-PD-1 treatment, and only 3 out of 12 tumours rebounded after treatment stopped. The survival rate of host mice bearing sgNipbl colon26 tumours was significantly improved after anti-PD-1 therapy compared to that of the vehicle group mice (p<0.0001, Fig. 3B). In contrast, anti-PD-1 therapy had little or no effect on tumour growth and the survival rate of mice bearing sgControl colon26 tumours compared to the vehicle group mice (p>0.05). Notably, anti-PD-1 therapy potently rejuvenated the CD8 TIL subpopulation compared to that of the vehicle group, as evidenced by increased Ki-67 and granzyme B expression levels (p<0.01, Figs. 3C and S4B). We obtained similar results in a CT26 murine syngeneic colorectal tumour model. As shown in Fig. 3D and 3E, 19 out of 28 *Nipbl*-deficient CT26 tumours in mice completely disappeared after anti-PD-1 treatment; the survival rate of host mice bearing sg*Nipbl* tumours was significantly prolonged compared to that of the vehicle group mice; In contrast, anti-PD-1 treatment had a weak inhibitory effect on sgControl CT26 tumours. Our data suggest that the PD-1/PD-L1 inhibitory checkpoint pathway is critical for maintaining the *in vivo* growth of *Nipbl*-deficient tumours.

**Figure 3.**
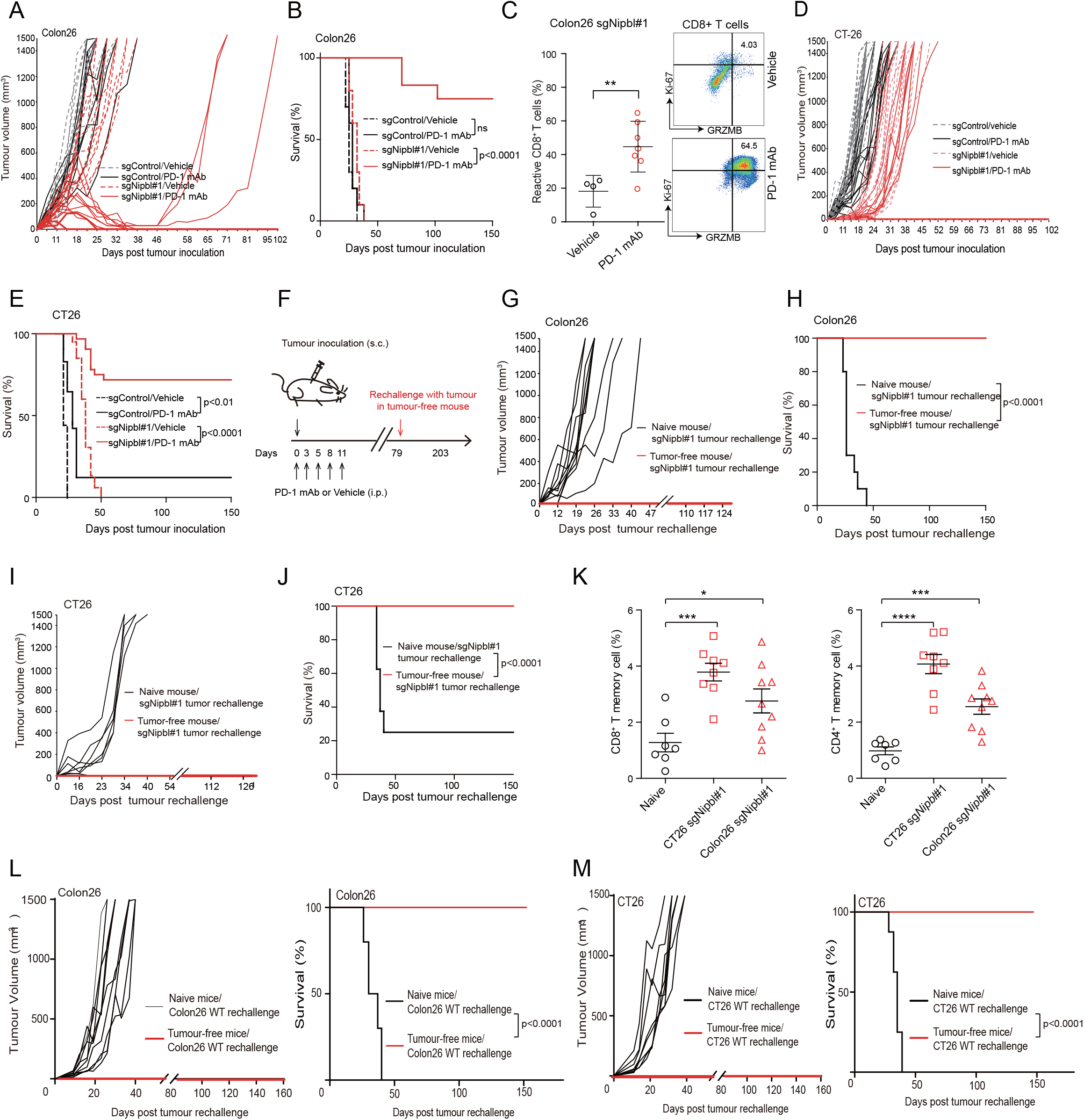
Nipbl-deficient tumours are supersensitive to anti-PD-1 therapy *in vivo*, eliciting antitumour memory responses in the host. (A-B) The therapeutic responses and survival rates of syngeneic mice bearing sgControl or sgNipbl colon26 tumours receiving anti-PD-1 mAb or vehicle treatment (sgControl group: vehicle, n=10; anti-PD-1, n=10; sgNipbl group: vehicle, n=10; anti-PD-1, n=12). (C) The percentages of reactive cytotoxic CD8+ T cells showing Ki-67+ and granzyme B+ expression in sgNipbl colon26 tumours after vehicle or PD-1 mAb treatment were quantitated (vehicle, n=4; anti-PD-1, n=7). Representative plots showing the gating for reactive CD8+ T cells showing Ki-67+ and granzyme B+ expression. (D-E) The therapeutic responses and survival rates of syngeneic mice bearing sgControl or sgNipbl CT26 tumours treated with anti-PD-1 mAb or vehicle (sgControl group: vehicle, n=18; anti-PD-1, n=18; sgNipbl group: vehicle, n=16; anti-PD-1, n=28). (F) Schematic diagram of the tumour re-challenge experiment. (G-H) Tumour growth and survival of tumour-free (n=8) and age-matched naïve mice (n=10) re-challenged with 1×10^6^ sgNipbl colon26 tumours 79 days after the first round of anti-PD-1 therapy. (I-J) Tumour growth and survival of tumour-free (n=16) and age-matched naïve mice (n=8) re-challenged with 5×10^5^ sgNipbl CT26 tumours 79 days after the first round of anti-PD-1 therapy. (K) The percentages of CD62L^+^/CD44^+^ central memory CD8+ T and CD4+ T cells in the spleens of tumour-free mice (colon26, n=9; CT26, n=8; age-matched naïve mice, n=7). (L) Tumour growth and survival of tumour-free (n=4) and age-matched naïve mice (n=10) re-challenged with 1×10^6^ parental colon26 tumours post anti-PD-1 therapy. (M) Tumour growth and survival of tumour-free (n=8) and age-matched naïve mice (n=8) re-challenged with 5×10^5^ parental CT26 tumours post anti-PD-1 therapy. The significance of differences in survival was assessed by the log-rank Mantel-Cox test. Data are shown as the mean ± SD, and the p value was calculated by two-tailed t tests. *p < 0.05, **p < 0.01, ***p < 0.001, ****p < 0.0001.

Next, we asked whether anti-PD-1 therapy elicits an antitumour memory response in the host (Fig. 3F). Tumour re-challenge data showed that *Nipbl*-deficient colon26 tumours were no longer able to grow in mice with completely regressed tumours even 124 days after tumour cell re-injection, but they developed quickly in age-matched naïve mice (Fig. 3G). The survival rate of these tumour-free mice was significantly prolonged compared to that of the naïve mice (p<0.0001, Fig. 3H). We obtained similar results in the CT26 mouse model (Fig. 3I and 3J). Flow cytometry analysis revealed that the proportions of both CD8+ and CD4+ central memory T cells were significantly increased in the spleens of tumour-free mice compared to those of naïve mice (p<0.05, Figs. 3K and S6), suggesting that anti-PD-1 therapy elicits antitumour immune memory responses in the host.

Because NIPBL is critical for maintaining genome instability^32^, *NIPBL*-defective tumours potentially increase their sensitivity to immunotherapy by producing neoantigens. To exclude this possibility, we individually transplanted *Nipbl* wild-type colon26 and CT26 parental tumours into sgNipbl tumour-free mice and observed that these tumours were again fully rejected compared to those of age-matched naïve mice, showing a significantly improved survival rate in the host (Fig. 3L and 3M), suggesting that *Nipbl* loss did not affect tumour cell immunity by introducing neoantigens in our case.

### *NIPBL* loss stimulates ERV expression in tumour cells by blocking DNMT1 transcription

To explore the mechanism by which *Nipbl* loss induces PD-L1 expression in tumours, we profiled the transcriptome of HCT-116 and CT26 cell lines with sgNIPBL, sgSTAG2, shSMC1A and their counterparts (Fig. 4A). The knockout and knockdown efficiencies of these subclones were shown in Fig. S3. We obtained 71 upregulated and 7 downregulated genes that overlapped in both HCT-116 and CT26 cell lines (average fold > 1.5). Metascape analysis^43^ revealed that genes related to type I/II interferon responses (GO: 0060337/0034341) and the host defence response to virus (R-HAS-1169410, hsa05168) were robustly clustered when comparing the cohesin loss and control groups (p<0.01, Fig. 4B). Protein-protein interaction analysis also revealed that the host defence response to virus network (GO: 0051607) including IRF9, BST2, IFIT1, RSAD2, IFIT3, OAS3 and STAT2 was significantly enriched when cohesin genes were silenced (in orange, Fig. 4A and 4C). Moreover, genes related to cellular response to exogenous dsRNA (GO: 0071360), such as DDX58/RIG-I, IFIH1/MDA5 and LGALS9, were highly enriched in the top 30 overlapping genes when comparing the sgNIPBL and sgControl groups (in red, Fig. 4A and 4B). Genes related to dsRNA binding (GO: 0003725), such as RIG-I, MDA5 and OAS3, were also enriched (Fig. 4A). As a positive control, the mRNA expression levels of PD-L1 (encoded by the CD274 gene, in blue) were significantly increased in all the tested samples. This observation led us to investigate whether dsRNA ERVs are actively expressed in *NIPBL*-deficient tumour cells.

**Figure 4.**
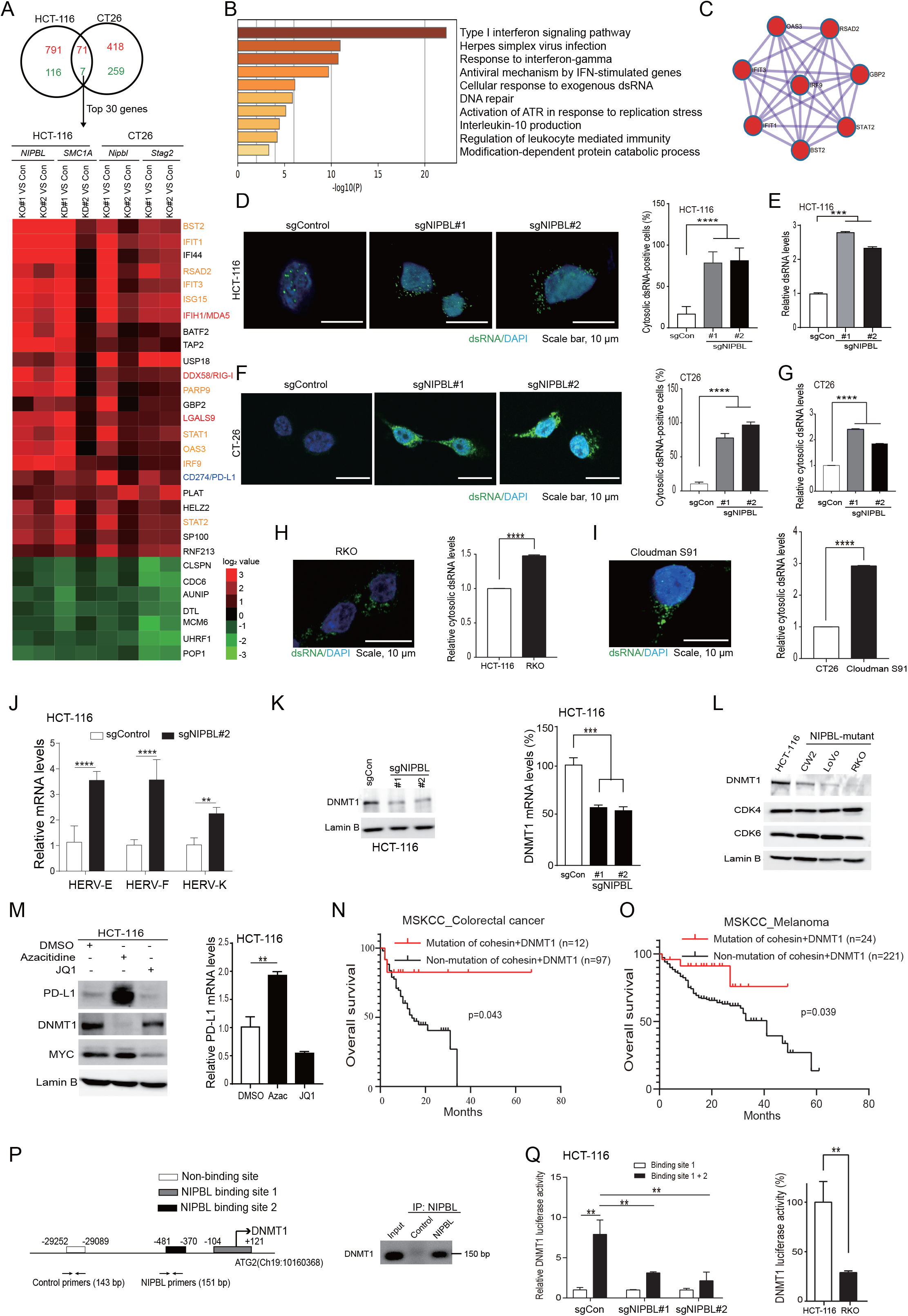
*NIPBL* loss stimulates ERV expression in tumour cells by blocking DNMT1 transcription. (A) The top 30 overlapping genes of the indicated groups were subjected to unbiased clustering by CLUSTER 3.0. (B) Representative enriched gene signatures by comparing the cohesin loss group to control groups. (C) IRF9-based protein interaction network was selected out by Metascape analysis. (D) dsRNA in HCT-116 cells was detected by a dsRNA-specific antibody using immunofluorescent staining. Bar diagram of cytosolic dsRNA-positive cells with statistics. (E) The dsRNA content was measured after removing DNA. (F) Immunofluorescent staining of dsRNA in sgControl and sgNipbl CT26 cells. Bar diagram of cytosolic dsRNA-positive cells with statistics. (G) The cytosolic dsRNA content was measured after removing DNA. (H-I) The cytosolic dsRNA of NIPBL-deficient RKO and cloudman S91 cells was imaged and quantified, while NIPBL wild-type HCT-116 and CT26 cells were used as controls. (J) The mRNA expression levels of HERV-E, HERV-F and HERV-K were determined by qRT-PCR. (K) The DNMT1 protein and mRNA expression levels were determined in sgControl and sgNIPBL HCT-116 cells. (L) The DNMT1, CDK4 and CDK6 protein levels were detected in three NIPBL-mutant colorectal tumour cell lines. (M) The protein and mRNA expression levels of the indicated genes were determined after treated with 2 μM azacitidine or JQ1 for 36 hours. (N) Mutations in STAG2, RAD21 and DNMT1 significantly prolonged the overall survival of colorectal cancer patients treated with anti-PD-1 therapy (n=12) compared with that of non-mutation counterparts (n=97) (log-rank p=0.043). (O) Mutations in STAG2, RAD21 and DNMT1 significantly prolonged the overall survival of melanoma patients treated with anti-PD-1 therapy (n=24) compared with that of non-mutation counterparts (n=221) (log-rank p=0.039). (P) ChIP-PCR analysis of NIPBL binding sites located in the DNMT1 promoter. (Q) The luciferase activity of the indicated DNMT1 promoter in HCT-116 sgControl, sgNIPBL and RKO cells. Data are shown as the mean ± SD, and the p value was calculated by two-tailed t tests. *p < 0.05, **p < 0.01, ***p < 0.001, ****p < 0.0001.

Immunofluorescent staining data showed that dsRNAs were dramatically increased in the cytosol of sgNIPBL HCT-116 cells, accounting for approximately 75% of total cells but were limited to the nucleus of sgControl cells (Fig. 4D). The dsRNA content in sgNIPBL HCT-116 cells was increased two-to-threefold compared to that of sgControl cells (Fig. 4E). Similarly, cytosolic dsRNAs were clearly observed at a higher level in sgNipbl CT26 cells compared to sgControl cells, accounting for approximately 75-90% of the total cells (Fig. 4F). The dsRNA content in the cytosol of sgNipbl CT26 cells was increased twofold compared to that of sgControl cells (Fig. 4G). dsRNAs were also clearly observed in the cytosol of endogenous *NIPBL*-deficient RKO and cloudman S91 tumour cell lines compared to wild-type HCT-116 and CT26 cells (Fig. 4H and 4I). The qRT-PCR data confirmed that human endogenous retroviruses (HERVs) such as HERV-E, HERV-F and HERV-K were significantly enhanced in sgNIPBL HCT-116 cells compared to sgControl cells (Fig. 4J). Based on these results, we conclude that *NIPBL* loss induces ERV expression in tumour cells.

ERVs are DNA sequences of retroviral origin that integrate into the mammalian genome, accounting for approximately 8-10% of the human and mouse genomes^44, 45^. The transcription of ERVs is strictly repressed in somatic cells by DNMT1-mediated DNA methylation to avoid the activation of host antiviral innate immunity^46, 47^. Thus, we asked whether *NIPBL* loss would affect DNMT1 expression. Expectedly, DNMT1 was markedly decreased at both the mRNA and protein levels when *NIPBL* was knocked out in HCT-116 cells (Fig. 4K). DNMT1 protein expression was also significantly reduced in three endogenous *NIPBL*-mutant human colorectal cancer cell lines (Fig. 4L). We did not observe obvious changes in either CDK4 or CDK6 protein levels in this case^48^. Next, we repeated this experiment using DNMT1 inhibitor 5’-azacytidine^49^. Compared to the vehicle, 5’-azacytidine robustly induced PD-L1 expression at both the mRNA and protein levels in HCT-116 cells (Fig. 4M). In contrast, the Myc inhibitor JQ1 had little or no influence on PD-L1 expression. This suggests that NIPBL loss-induced DNMT1 inhibition but not Myc inhibition is critical for inducing PD-L1 expression in tumour cells^50^.

Noteworthy, anti-PD-1 therapy significantly prolonged overall survival of colorectal cancer and melanoma patients harbouring mutations in *STAG2*, *RAD21* and *DNMT1* compared to that of their counterparts without mutations (colorectal cancer: p=0.043; melanoma: p=0.039, Fig. 4N, 4O and Supplemental Table 1)^51^. Because only the genetic mutations of STAG2 and RAD21 but no other members of cohesin core subunits and regulators are assessable in these Memorial Sloan Kettering Cancer Center (MSKCC) immunogenomic studies^51^, we included DNMT1 mutation based on the mechanism of cohesin loss-induced DNMT1 inhibition.

We next asked whether NIPBL directly regulates DNMT1 transcription. By assessing the ENCODE data^52^, we identified that NIPBL may interact with the DNMT1 promoter. The chromatin immunoprecipitation (ChIP)-PCR data showed that NIPBL selectively bound with the DNMT1 promoter in the -481-bp to +121-bp region compared to that of the control primer (Fig. 4P). We obtained similar results using IgG as a negative control. There were two potential NIPBL binding sites located in the -481-bp to +121-bp region of the DNMT1 promoter (Fig. 4P, left)^52^. Thus, we individually cloned the DNMT1 promoter (-481 bp to +121 bp and -269 bp to +121 bp) into a pGL3-basic luciferase vector and observed that the luciferase activity of the DNMT1 promoter (-481 bp to +121 bp) was greatly impaired when binding site 2 (-481 bp to -370 bp) was removed (Fig. 4Q, left). The luciferase activity of the DNMT1 promoter (-481 bp to +121 bp) was significantly reduced in sgNIPBL HCT-116 cells compared to sgControl cells. Similarly, the luciferase activity of the DNMT1 promoter (-481 bp to +121 bp) was also impaired in *NIPBL*-deficient RKO cells compared to wild-type HCT-116 cells (Fig. 4Q, right). This suggests that the binding site 2 is critical for NIPBL-mediated DNMT1 transcription. Collectively, we conclude that *NIPBL* loss provokes ERV expression by impairing DNMT1 transcription.

### The dsRNA-RIG-I/MDA5-MAVS-IRF3 signalling is responsible for *NIPBL* loss-induced PD-L1 expression

The RIG-I/MDA5-MAVS signalling pathway is responsible for detecting cytosolic viral RNA^53–58^. Immunoblotting data showed that the core members of the RIG-I/MDA5-MAVS signalling machinery, such as RIG-I, MDA5, MAVS, IRF3 and IRF7, were robustly increased when *NIPBL* was knocked out in HCT-116 cells (Fig. 5A). We also observed that STAT1, STAT2 and IRF9, which are key regulators for type I/II interferon responses^59^, were highly expressed in sgNIPBL HCT-116 tumour cells compared to sgControl cells. Similar results were obtained in *NIPBL*-mutant CW2 and LoVo colorectal tumour cell lines, which showed increased PD-L1 expression compared to that in control cells (Fig. 5B). To investigate whether RIG-I/MDA5-MAVS signalling is required for *NIPBL* loss-induced PD-L1 expression, we silenced MAVS using three different shRNAs and observed that shMAVS obviously reduced PD-L1 expression in sgNIPBL HCT-116 cells compared to scramble cells (Fig. 5C). MAVS knockdown also greatly reduced PD-L1 expression in *NIPBL*-mutant RKO tumour cells (Fig. 5D). Notably, exogenous overexpression of wild-type MAVS recaptured the transcriptional signature of the sgNIPBL group, which (compared to the control group) showed increased mRNA and protein expression of genes related to the defence response to virus (GO: 0051607), such as RIG-I, MDA5, STAT1, STAT2 and IRF9 plus CD274 (Fig. 5E and 5F). We did not observe a significant change in IRF3 protein due to the large amount of IRF3 protein in 293T cells, but the phosphorylation of IRF3 at serine 386 was increased. This suggests that the dsRNA-MAVS signalling is crucial for the *NIPBL* loss-induced cell phenotype.

**Figure 5.**
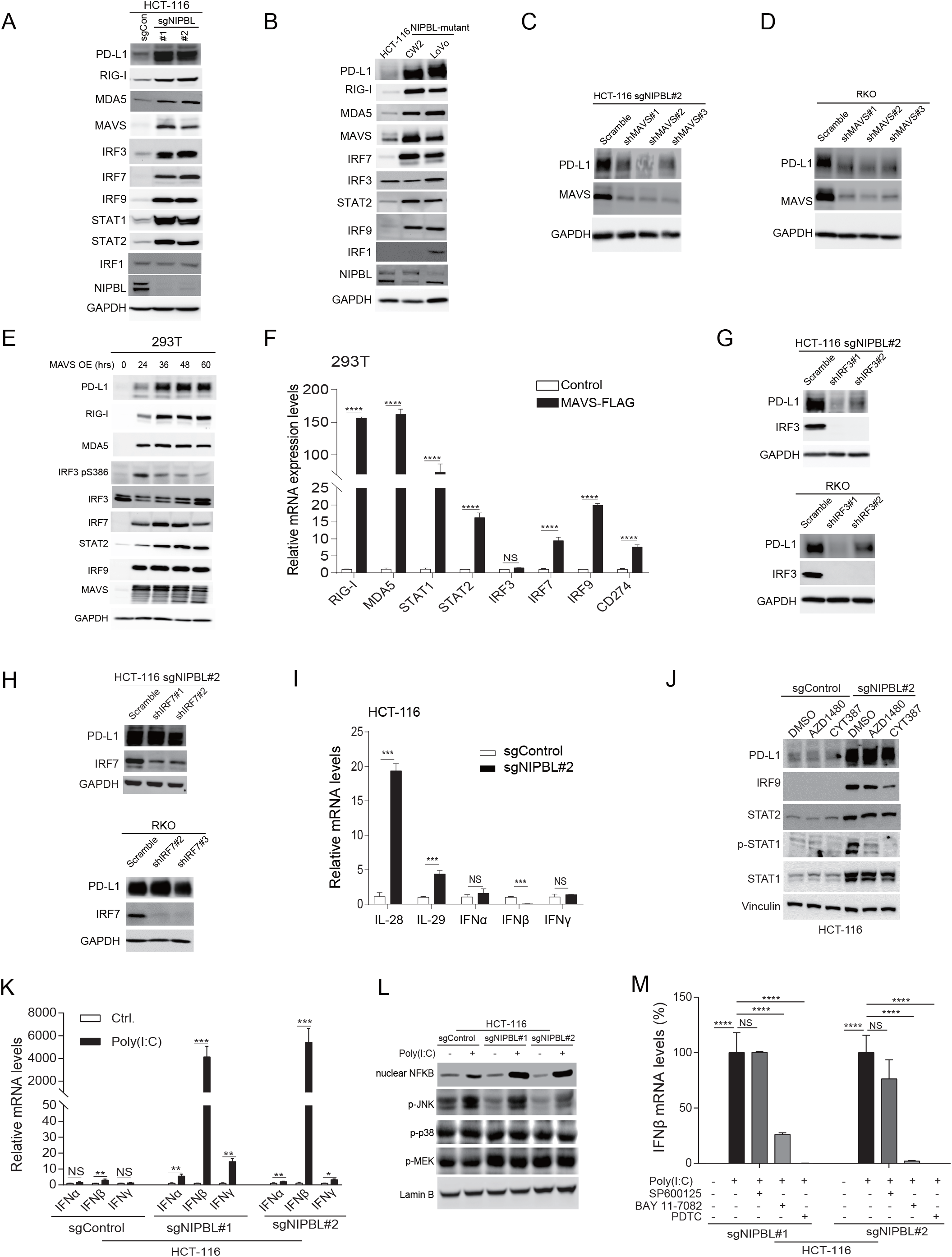
The dsRNA-MAVS-IRF3 signalling is responsible for *NIPBL* loss-induced PD-L1 expression. (A-B) The indicated proteins of the dsRNA-MAVS signalling machinery plus other proteins were detected by immunoblotting. (C-D) PD-L1 protein expression was detected by immunoblotting when MAVS was knocked down. (E-F) The mRNA and protein levels of the dsRNA-MAVS signalling machinery were detected when MAVS was exogenously overexpressed. (G) PD-L1 protein expression was detected in sgNIPBL HCT-116 and RKO cells by immunoblotting when IRF3 was knocked down. (H) PD-L1 protein expression was detected in sgNIPBL HCT-116 and RKO cells by immunoblotting when IRF7 was knocked down. (I) The mRNA expression levels of IFN-α, IFN-β, IFN-γ, IL-28 and IL-29 were determined in sgControl and sgNIPBL HCT-116 cells by qRT-PCR. (J) The indicated protein expression levels were determined by immunoblotting in the absence or presence of 0.5 μ M AZD1480 or CYT387. (K) The mRNA expression levels of IFN-α, IFN-β and IFN-γ were determined in sgControl and sgNIPBL HCT-116 cells in the absence or presence of 500 ng/ml poly (I:C) treatment. (L) Indicated proteins were detected by immunoblotting in the absence or presence of 500 ng/ml poly (I:C) treatment. (M) The IFN-β mRNA level was determined by qRT-PCR with or without NF-κ B inhibitor (BAY 11-7082 and PDTC) or JNK inhibitor (SP600125) treatment. Data are shown as the mean ± SD, and the p value was calculated by two-tailed t tests. *p < 0.05, **p < 0.01, ***p < 0.001, ****p < 0.0001.

IRF3 and IRF7 are two major downstream effectors of the dsRNA-MAVS signalling^60, 61^, and both showed increased protein expression levels in *NIPBL*-deficient tumour cells compared to control cells (Fig. 5A and 5B). Thus, we assessed whether IRF3 or IRF7 is critical for *NIPBL* loss-induced PD-L1 expression. shRNA silencing data demonstrated that IRF3 knockdown abrogated PD-L1 protein expression in *NIPBL*-deficient HCT-116 and RKO tumour cells (Fig. 5G). In contrast, IRF7 knockdown had little or no effect on PD-L1 expression (Fig. 5H). These data suggest that the MAVS-IRF3 signalling is critical for *NIPBL* loss-induced PD-L1 expression.

In the literature, the activation of MAVS-IRF3 signalling often leads to the production of IFN-β in non-immune cells, which is critical for antiviral innate immunity^54–58^. The qRT-PCR data revealed that the mRNA expression levels of IFN-α,IFN-β and IFN-γ , but not type III interferons IL-28 and IL-29, were low or undetectable in *NIPBL*-deficient tumour cells (Fig. 5I), which is consistent with previous reports^61, 62^. We also detected no phosphorylation of STAT2 at Tyr 690 in sgNIPBL tumour cells, which is rapidly activated by JAK upon IFNs stimulation^59^. We further treated sgNIPBL HCT-116 cells with two JAK inhibitors and observed that both AZD1480 and CYT387 had no significant effect on PD-L1 protein expression by comparing sgNIPBL HCT-116 cells and sgControl cells, although STAT1 Y701 phosphorylation was largely abolished (Fig. 5J). In addition, we individually incubated sgControl HCT-116 and colon26 cells with sgNIPBL tumour cell medium for 48 hours and observed no change in PD-L1 protein expression (Fig. S7). These data suggest that NIPBL loss-induced PD-L1 expression is not IFNs-dependent.

Using dsRNA synthetic analog polyinosinic-polycytidylic acid (poly (I:C)) as a positive stimuli, we observed that IFN-β but not IFN-α/γ was robustly increased approximately 4000-fold in sgNIPBL HCT-116 cells compared to that of sgControl group (Fig. 5K). Because the IFN-β transcription is co-ordinately regulated by the IFN-β enhanceosome consisting of IRF3/7, NF-κB and phosphorylated JNK, MEK and p38-mediated AP-1 activation ^63, 64^, we profiled these protein expression levels and observed that nuclear NF-κ B and phosphorylated JNK were weakly expressed in sgNIPBL HCT-116 cells, but were highly activated by poly (I:C) treatment compared to that of vehicle (Fig. 5L). p38 and MEK remained highly phosphorylated in all the cases. To examine whether NF-κB or JNK activation is crucial for IFN-β expression, we individually treated sgNIPBL HCT-116 cells with small molecular inhibitors of NF-κB (BAY 11-7082 and pyrrolidinedithiocarbamate ammonium (PDTC)) or JNK (SP600125) and observed that two NF-κB inhibitors but not JNK inhibitor dramatically reduced approximately 75-95% of poly (I:C)-induced IFN-β expression in sgNIPBL HCT-116 cells (Fig. 5M), suggesting that NF-κB activation is essential for IFN-β expression. Noteworthy, NIPBL-induced MAVS-IRF3/7 activation is also crucial for IFN-β expression because sgControl HCT-116 tumour cells with lowly expressed IRF3/7 had a very weak IFN-β expression upon poly (I:C) treatment. Taken together, we conclude that NIPBL loss-induced ERV expression selectively increase PD-L1 expression but not IFN-β due to lack of NF-κB activation.

### IRF3 stimulates PD-L1 expression by assembling the STAT2-IRF9 complex

To further explore the mechanism of PD-L1 regulation, we cloned the PD-L1 promoter into a pGL3-basic vector and observed that the luciferase activity of the PD-L1 promoter (the -799-bp to +153-bp region from the translational starting site) was 50-fold higher than that of the vector control (Fig. 6A), suggesting that this promoter region is critical for PD-L1 transcription. Promoter truncation analysis revealed that the luciferase activity of the PD-L1 promoter was reduced by 80% in sgNIPBL HCT-116 cells when the -223-bp to -133-bp region was removed (Fig. 6B). We obtained similar results in *NIPBL*-mutant RKO cells (Fig. 6C). This suggests that the -223-bp to -133-bp region of the PD-L1 promoter is responsible for *NIPBL* loss-induced PD-L1 expression. DNA binding consensus motif analysis revealed that one STAT2 and one IRF binding site were potentially located in this small region (Fig. 6A). As previously mentioned, STAT2, IRF3, IRF7 and IRF9 were markedly increased in NIPBL-deficient tumour cells compared to control cells, while IRF1 was weakly or not expressed (Figs. 5A, 5B and 6D). Thus, we asked whether IRFs or its combination with STAT2 could induce PD-L1 expression. Using IRF1 as a positive control^65^, exogenous overexpression of IRF1, IRF1 plus STAT2 and STAT2 plus IRF9 robustly enhanced the luciferase activity of the 386-bp PD-L1 promoter, while STAT2 alone, other IRFs or their combination with STAT2 failed to do so (Fig. 6E). Remarkably, individual knockout of STAT2 and IRF9 obviously abrogated PD-L1 expression in sgNIPBL HCT-116 cells compared to that in the control cells (Fig. 6F). In contrast, IRF1 knockdown had little or no influence on PD-L1 expression (Fig. 6G). Knockdown of STAT1, which was located outside of the -223-bp to -133-bp region, also had no effect on PD-L1 expression (Fig. 6H). The analysis of TCGA tumour data revealed that both STAT2 and IRF9 well correlated with PD-L1 expression at mRNA levels in stomach and colorectal tumour samples (Figs. 6I and S8). These results indicate that STAT2 and IRF9 are potentially responsible for NIPBL loss-induced PD-L1 expression.

**Figure 6.**
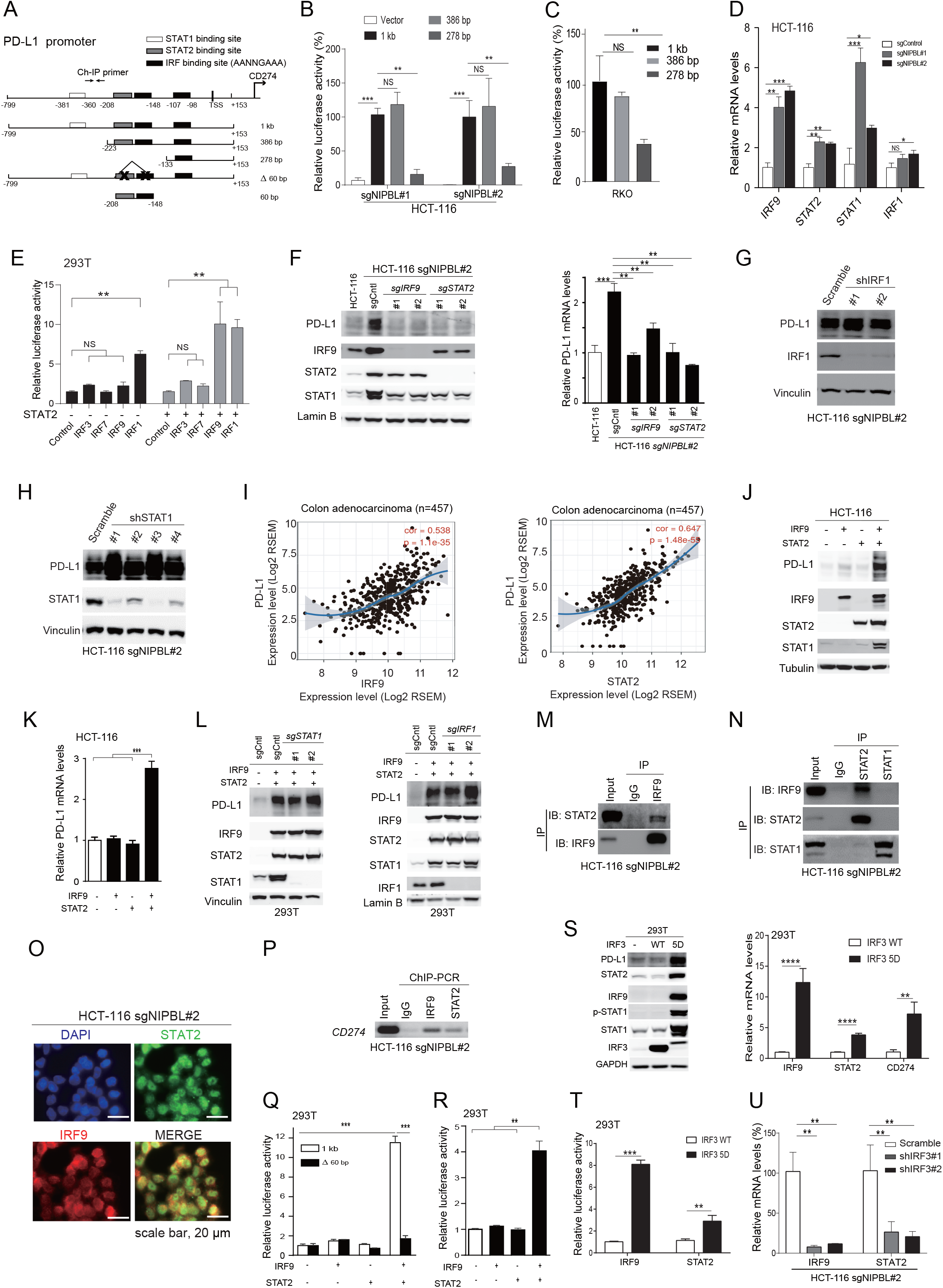
STAT2 and IRF9 co-regulate PD-L1 transcription independent of STAT1 and IRF1. (A) Schematic diagram of the PD-L1 promoter. (B-C) The relative firefly luciferase expression levels of the indicated PD-L1 promoter in sgNIPBL HCT-116 and RKO cells. Renilla luciferase was used as an internal control. (D) The mRNA expression levels of the indicated genes in sgNIPBL HCT-116 cells were determined by qRT-PCR. GAPDH was used as an internal control. (E) The luciferase activity of the 386-bp PD-L1 promoter was measured when IRFs alone or IRFs and STAT2 were overexpressed. Renilla luciferase was used as an internal control. (F) The indicated protein and mRNA expression levels were determined in sgControl, sgSTAT2 and sgIRF9 HCT-116 cells. (G) The indicated protein levels were determined by immunoblotting when IRF1 was knocked down in sgNIPBL HCT-116 cells. (H) The indicated protein levels were determined by immunoblotting when STAT1 was knocked down in sgNIPBL HCT-116 cells. (I) Correlation of STAT2 and IRF9 mRNA levels with PD-L1 mRNA levels in TCGA colorectal tumour samples (n=457). Spearman’s correlation and estimated significance were calculated with TIMER and adjusted for tumour purity. (J-K) The expression levels of the indicated protein and mRNA were determined by immunoblotting when STAT2 and IRF9 were co-expressed in HCT-116 cells. (L) The expression levels of the indicated protein and mRNA were determined by immunoblotting when STAT2 and IRF9 were co-expressed in sgSTAT1 or sgIRF1 293T cells. (M) STAT2 physically associated with IRF9. IgG was used as a negative control. (N) IRF9 selectively bound with STAT2 but not STAT1. (O) Dual immunofluorescent staining of STAT2 and IRF9 in sgNIPBL HCT-116 cells. Scale bar, 20 μm. (P) STAT2 and IRF9 co-occupied the PD-L1 promoter close to the -233-bp to -133-bp region in sgNIPBL HCT-116 cells according to a ChIP-PCR assay. (Q) The luciferase activity of the 1-kb PD-L1 promoter with or without the 60-bp STAT2-IRF9 binding site was detected when STAT2 and IRF9 were individually expressed or co-expressed. (R) The luciferase activity of the 60-bp PD-L1 promoter was significantly elevated when STAT2 and IRF9 were co-expressed. (S) Indicated protein and mRNA expression were determined when IRF3 wild-type and 5D constitutive active mutant were exogenously expressed for 48 hours. (T) The luciferase activities of STAT2 and IRF9 promoters were determined when IRF3 wild-type and 5D mutant were exogenously expressed for 48 hours. (U) The mRNA expression levels of STAT2 and IRF9 were determined when IRF3 was knocked down. Data are shown as the mean ± SD, and the p value was calculated by two-tailed t tests. *p < 0.05, **p < 0.01, ***p < 0.001, ****p < 0.0001.

Next, we asked whether co-expression of STAT2 and IRF9 induces PD-L1 expression. As expected, STAT2 and IRF9 robustly increased PD-L1 at both the mRNA and protein levels, accompanied by elevated STAT1 expression (Fig. 6J and 6K). To further exclude STAT1 and IRF1, we individually established sgSTAT1 and sgIRF1 293T clones and observed that co-expression of STAT2 and IRF9 robustly increased PD-L1 expression in these cells (Fig. 6L), suggesting that STAT2 and IRF9 co-ordinately regulate PD-L1 transcription independent of IRF1 and STAT1. In the literature, STAT2 but not STAT1 is evolutionarily conserved to interact with IRF9^66^. Thus, we performed an immunoprecipitation (IP) assay and observed that STAT2 physically associated with IRF9 in sgNIPBL HCT-116 cells (Fig. 6M). Using a reciprocal IP assay, we observed that IRF9 selectively binds with STAT2 but not STAT1 (Fig. 6N). Dual immunofluorescent staining data illustrated that STAT2 and IRF9 were largely co-localized in the nucleus of sgNIPBL HCT-116 cells (Fig. 6O).

The ChIP-PCR data confirmed that STAT2 and IRF9 co-occupied the PD-L1 promoter close to the -208-bp to -148-bp regions upstream of the translation start site (Fig. 6A and 6P). Luciferase expression data showed that co-expression of STAT2 and IRF9 increased PD-L1 promoter activity by approximately 11-fold, but this effect was completely abolished when the 60-bp STAT2-IRF9 DNA binding motif was removed (Fig. 6A and 6Q). We cloned this 60-bp DNA fragment and observed that it responded to the co-expression of STAT2 and IRF9 (Fig. 6R). Thus, we conclude that the STAT2-IRF9 complex is essential for *NIPBL* loss-induced PD-L1 expression in tumour cells.

Finally, we asked whether IRF3 is responsible for STAT2 and IRF9 expression. We generated a constitutive active IRF3 5D mutant by substituting Ser396, Ser398, Ser402, Thr404 and Ser405 with aspartic acids^67^ and observed that exogenously expression of IRF3 5D mutant but not wild-type robustly activated STAT2, IRF9 and PD-L1 at both mRNA and protein levels (Fig. 6S). Consistently, IRF3 5D mutant significantly increased the luciferase expression levels of IRF9 and STAT2 promoters containing the AANNGAAA motif compared to that of wild-type IRF3 (Fig. 6T). Meanwhile, shIRF3 markedly reduced the mRNA expression levels of both STAT2 and IRF9 in sgNIPBL HCT-116 cells compared to scramble (Fig. 6U). These results indicate that IRF3 promotes PD-L1 transcription in NIPBL-deficient tumours by transcriptionally upregulating the STAT2-IRF9 complex.

## Discussion

ERV expression, which often leads to immunosuppression in the host^68^, has recently been identified in numerous human cancer types and is well correlated with patient responses to immunotherapy^69, 70^. Our study reveals that ERV expression is a consequence of cohesin loss-mediated DNMT1 inhibition in solid tumour cells and inactivates host CD8 TILs via the PD-L1/PD-1 inhibitory checkpoint pathway. Mechanistically, ERV expression selectively promotes the PD-L1 expression by stimulating the dsRNA-MAVS-IRF3-STAT2/IRF9 signalling pathway, but doesn’t induce IFN-β expression due to lack of NF-κ B activation. *Villin*-driven *Nipbl* deficiency significantly promotes tumour growth in the host when crossed with *APC^min^* mice, reinforcing that *Nipbl*-deficient tumours are immunosuppressive. Noteworthy, cohesin loss leads to defective interferon responses in human acute myeloid leukaemia, of which high STAT1 and STAT2 expression levels are critical for the development and antiviral and antitumour immunity of immune cells^71–74^.

Nevertheless, our study provide a novel mechanism by which tumour-associated ERVs establish an immunosuppressive microenvironment.

Although DNMT1 inhibitors are clinically used for treating lymphomas by producing viral mimicry responses^61, 62^, mice with highly reduced Dnmt1 expression levels spontaneously developed T cell lymphoma^47^. The TCGA data analysis revealed that patients with low DNMT1 mRNA expression had a significantly worse survival rate than those with high DNMT1 expression in both stomach and colorectal tumour cohorts (Fig. S9). This observation led us to investigate whether PD-L1 is critical for the relapse of low DNMT1-expressing tumours. To verify this, we pretreated lewis lung cancer cells with 0.5 μM decitabine for 3 days and inoculated them into the flanks of syngeneic mice and observed that anti-PD-1 monoclonal antibody potently reduced the *in vivo* growth of decitabine-pretreated lewis lung cancer cells compared to that with vehicle, and 2 out of 3 tumours completely regressed (p=0.011, Fig. S10A). PD-L1 protein expression was visibly increased after treatment with decitabine (Fig. S10B). As a control, lewis lung cancer cells showed no significant difference between the vehicle and anti-PD-1 groups (p>0.05, Fig. S10C).

Consistently with previous report^75^, the PD-L1 expression was often increased after DNMT1 inhibitor treatment (Fig. 4M). Thus, it would be ideal to combine with anti-PD-1 therapy when applying DNMT1 inhibitors to patients when it elicits PD-L1 expression in tumour cells.

In the literature, it is generally accepted that a high mutation burden in tumours is associated with the clinical outcomes of anti-PD-1 therapy^31, 51, 76, 77^. Changes leading to deficiency of DNA mismatch repair, such as *Mlh1* loss, can increase the mutational burden of tumour cells and increase tumour sensitivity to immune checkpoint inhibitors^4, 78^. To assess whether mutations of cohesin subunits and regulators are correlated with high mutation load, we analysed MSKCC immunogenomic data and observed that mutations of STAG2, RAD21 and DNMT1 were well correlated with a higher mutation load in MSI-subtype colorectal cancer samples (n=109, p=4.42E-5, Supplemental Table 1); however, mutations of STAG2, RAD21 and DNMT1 were poorly associated with a high tumour mutation load in microsatellite stable (MSS) melanoma (n=245, p=0.92, Supplemental Table 1). The average mutation counts for melanoma and colorectal cancer were 773.1 and 27.7, respectively.

Because *NIPBL* is frequently mutated at repetitive DNA sequence sites, which is a characteristic of tumours with MSI^79–81^, we further generated *Mlh1* knockout CT26 cells and performed another round of GeCKO screening, showing that sgRNAs targeting cohesin subunits (*Stag1*, *Smc1b* and *Rad21*) and regulators (*Nipbl*, *Pds5a*, *Pds5b* and *Esco1)* were again selected out in the anti-PD-1/vehicle group compared to nontargeting sgRNA group (p<0.005, Fig. S11). As shown in Fig. 2E, murine B16F10 melanoma cells used in previous CRISPR-Cas9 library screens are Nipbl-deficient, which explains why cohesin genes were not profiled^22^. Notably, *Nipbl* depletion in gastrointestinal tissue is not sufficient to elicit tumour in mice, showing a normal lifespan and pregnancy in the host, suggesting that other cancer driver mutations are required to promote tumour formation *in vivo*. These results indicate that cohesin loss-induced tumour immunity is likely independent of a high tumour mutation load but has a greater chance of occurring when there is a high number of gene mutations.

As core components of the transcriptional machinery, DNA polymerases such as POLE and POLD1 are highly mutated in human cancer and are well associated with a high tumour mutation burden^31^. Thus, it is particularly of interest to explore whether DNMT1 inhibition commonly co-occurs with the deregulation of transcriptional machinery, including cohesin, mediators and DNA polymerases. If that were the case, low DNMT1 expression or ERV expression could serve as an independent biomarker for predicting the therapeutic responses to anti-PD-1 therapy.

In the literature, IFN-γ is well-known for boosting PD-L1 expression in tumour cells through STAT1-IRF1 signalling^65, 82, 83^. We observed little or no IFN-γ and IRF1 expression in *NIPBL*-deficient tumour cells, but STAT1 was actively expressed when ERV or IRF3 5D mutant was expressed (Figs. 5A and 6S). Individual knockdown of IRF1 and STAT1 did not affect PD-L1 expression in *NIPBL*-deficient tumour cells. JAK inhibitor also didn’t affect PD-L1 expression, although it potently blocked STAT1 phosphorylation at Tyr701 site. These results suggest that the STAT1-IRF1 signalling is dispensable for ERV-induced PD-L1 expression. Interestingly, depletion of STAT2 and IRF9 had also no impact on IFN-γ-induced PD-L1 expression compared to that with STAT1 knockout (Fig. S12). These results indicate that the STAT2-IRF9 complex plays a unique role in ERV-induced PD-L1 expression, which is separate from IFN-γ -induced PD-L1 expression via the STAT1-IRF1 complex.

In summary, our study reveals a novel mechanism by which *NIPBL* deficiency regulates immunity in cancer by stimulating ERV expression; *NIPBL*-deficient tumours are highly immunosuppressive, but this effect could be rejuvenated by anti-PD-1 therapy.

## Material and methods

### Cell lines

CT26, Lewis lung cancer, B16F10, HEK-293T, CW2, LoVo, RKO and HCT-116 cell lines were purchased from the Cell Bank of Shanghai Institutes for Biological Sciences, Chinese Academy of Sciences (Shanghai, China). Colon26 and cloudman S91 murine cell line was purchased from BeNa Culture Collection (Beijing, China). All cells were maintained in DMEM/F12 or RPMI 1640 medium supplemented with 10% fetal bovine serum and 1% penicillin streptomycin and incubated at 37°C in a humidified incubator with 5% CO_2_. All human cell lines were authenticated by STR. The strain source of mouse cell lines was identified using PCR^84^. CT26 and colon26 were subjected to exon sequencing. All cell lines were tested negative for mycoplasma contamination.

### *In vivo* mouse GeCKO screening

The GeCKO screening was performed following previously reports^19, 85, 86^. CT26 cells were infected with lentiviruses containing GeCKO library sgRNAs at a multiplicity of infection (MOI) of 0.3, and were selected with puromycin for 7 days to remove uninfected cells. After selection, 2×10^7^ cells were directly harvested as DMSO_Day0_ group and frozen for genomic DNA extraction, and the remaining cells were inoculated into the flanks of syngeneic mouse for receiving i*n vivo* screening (2×10^7^ cells per injection, 10 mice per group). Mice were randomly assigned to two groups treated with control IgG or anti-PD-1 monoclonal antibody (ip. 200 μg/mouse, clone RMP1-14, Bio X cell) at day 0, 3, 5, 8, 11 post inoculation, respectively.

After 21 days post inoculation, the genomic DNA was individually extracted from tumour tissues of each mouse and was amplified for Illumina sequencing using two-step PCR^50^. The second PCR products were purified using Qiagen gel extraction kit. To avoid tumour size imbalance, equal amounts of PCR products of each tumour sample was picked up and mixed in each group for Illumina HiSeq X10 sequencing by Novogen according to the manufacturer’s instruction. The sgRNA averages of DMSO_day0_, Control_day21_ and anti-PD-1_day21_ groups were 487, 404 and 516, respectively, which was approximately 400-fold of total library sgRNAs. The library coverage rates of DMSO_day0_, Control_day21_ and anti-PD-1_day21_ groups were 99.35% (65531/65959 sgRNAs), 68.06% (44894/65959 sgRNAs) and 89.68% (59156/65959 sgRNAs), respectively. The loss of library sgRNAs in the Control_day21_ group is likely caused by tumour overgrowth.

After normalized with medium normalization, P-values of sgRNAs were calculated using a MAGeCK algorithm that test whether sgRNA abundance differs significantly between control IgG and anti-PD-1 groups. sgRNAs were then ranked based on P-values using a modified robust ranking aggregation algorithm to identify positively or negatively selected genes^24^. The fold changes of sgRNAs were calculated by comparing the read counts of anti-PD-1 group to that of control IgG group after normalized. Similar experimental procedure was applied in *Mlh1* knockout CT26 tumour screening.

### *In vivo* anti-PD-1 treatment

All the mice were purchased from Shanghai SLAC Laboratory Animal Co. Ltd. All studies were performed according to the guidelines of Institutional Animal Care and Use Committee of Shanghai Institute of Nutrition and Health. For xenograft tumours, 5×10^5^ CT26 or 1×10^6^ colon26 cells were subcutaneously injected into the flanks of the 6-week-old syngeneic BALB/C mice. After injected, mice were randomly assigned to different groups, followed by the treatment of vehicle or anti-PD-1 antibody (i.p., 200 μg/mouse, Bio X cell, clone RMP1-14) at day 0, 3, 5, 8, 11 post inoculation. Tumour size was measured twice one week, and tumour volume was determined based on the following formula: Tumour volume = 0.5 × length × width^2^.

### Statistics

All statistical analyses were done in GraphPad Prism 7 software. Survival significance was assessed by a Logrank Mantel-Cox test. All data were presented as means ± SD. The p value was calculated by two-tailed *t* tests. *, *p* <0.05, **, *p* < 0.01, ***, *p* < 0.001.

### Data Availability

All the datasets are available from the corresponding author upon reasonable request. sgRNA library screening sequencing data and RNA-Seq data have been deposited into NCBI database under accession number GSE125201, GSE125202 and GSE124909.

Please see more information in supplemental data.

## Acknowledgments

This work was financially supported by the CAS_Key Research Program of Frontier Sciences (QYZDB-SSW-SMC034), Strategic Priority Research Program of Chinese Academy of Sciences (XDA12020210), National Natural Science Foundation of China (81322034) and National Key R&D Program of China (2016YFC1302400). Thanks for the technical support of Ms. Yongfeng Zhang, Institutional Core facility.

## Author contributions

J.Y.L conceived and supervised the study. J.Y.L and Y.H. wrote the manuscript. Y.H. performed the experiments on sgRNA library screening, Nipbl knockout cell line, in vivo anti-PD-1 therapy and STAT2-IRF9 function. F. P. performed the dsRNA- and DNMT1-related experiments. Y.C. performed the experiments on the MAVS-IRF3 signalling and Nipbl knockout mice. T. L. performed experiments on NIPBL^fl/fl^; APC^Min^ mice, Nipbl knockout cell line and statistical analysis on mutations of cohesin subunits and regulators in TCGA samples. J.S. performed Nipbl wild-type tumour re-challenge experiments. Z.C. performed the statistical analysis of melanoma and colorectal cancer patients responding to anti-PD-1 therapy. Q.D helped on flow cytometric analysis of tumour infiltrated immune cells. F.J., H.X., H.Z., X.K., Q.C., J.D. contributed to discussion of results.

