## supplemental data for "Cohesin mutation sensitizes cancer cells to anti-PD-1 therapy through endogenous retrovirus-mediated PD-L1 upregulation"

Supplemental Figures

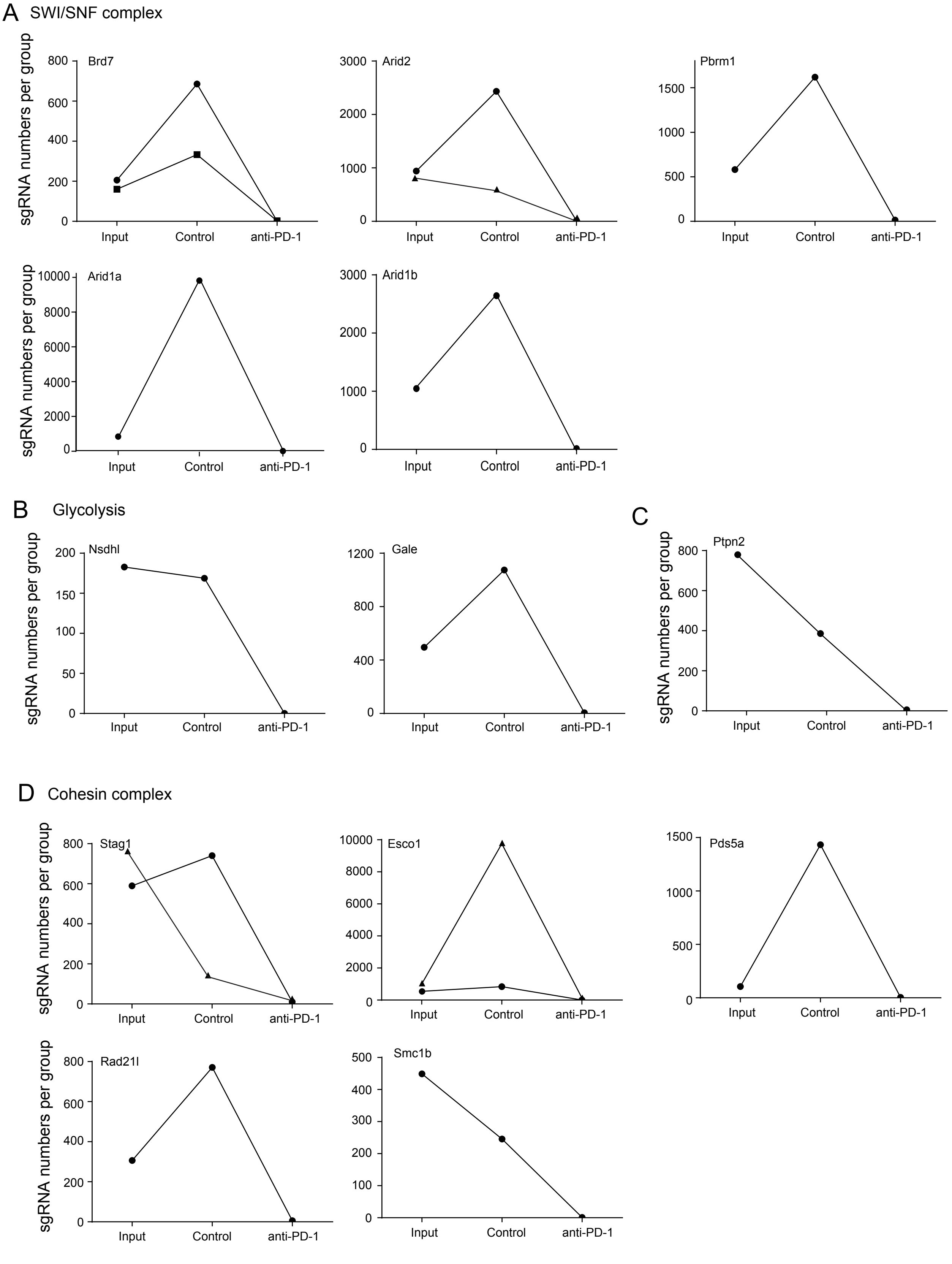


**Supplemental Figure 1.** The sgRNAs targeting SWI/SNF complex genes (A), glycolysis-related genes (B), Ptpn2 (C) and cohesin subunits and regulators (A) were significantly reduced in anti-PD-1 group by comparing to control IgG group. sgRNAs in the $DMSO_{day0}$ group was used as input after normalization.

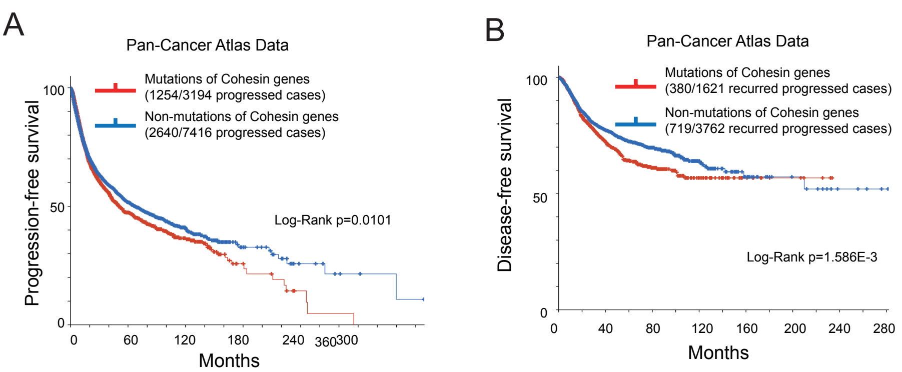


**Supplemental Figure 2.** Mutations of cohesin subunits and regulators negatively correlated with the patients' progression-free (A) and disease-free (B) survivals. TCGA data was obtained from cbioportal website.

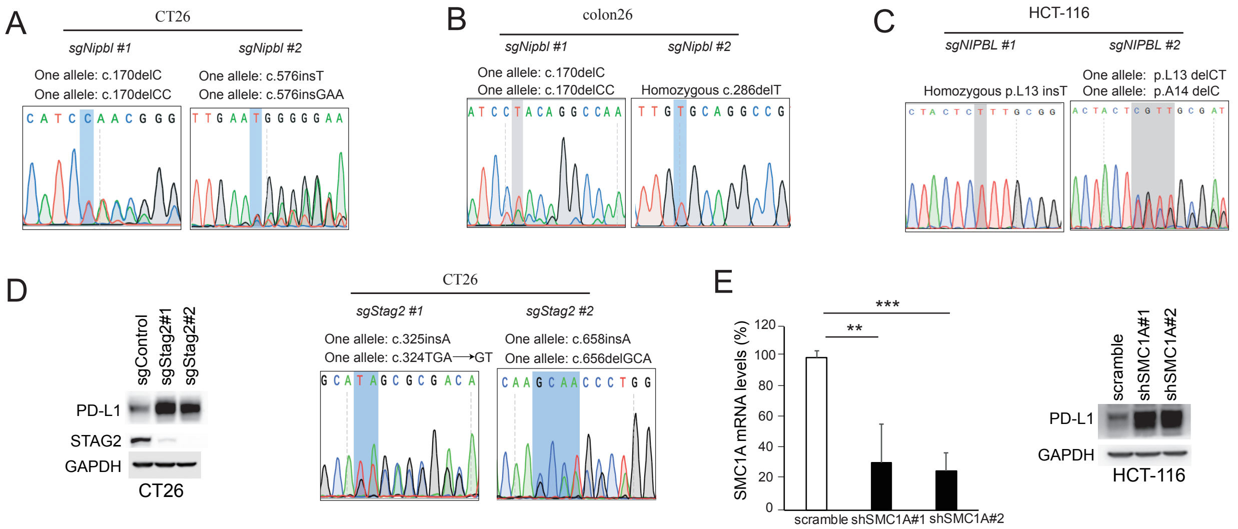


**Supplemental Figure 3.** Validation of *Nipbl* knockout in CT26, colon26 and HCT-116 cells. (A) Validation of *Nipbl* knockout (mouse sgNipbl targeting sequence #1 and #2) in CT26 cells using Sanger sequencing. (B) Validation of *Nipbl* knockout (mouse sgNipbl targeting sequence #1 and #3) in colon26 cells using Sanger sequencing. (C) Validation of *NIPBL* knockout in HCT-116 cells using Sanger sequencing. (D) Validation of *Stag2* knockout in CT26 cells using immunoblotting and Sanger sequencing. (E) Validation of SMC1A knock-down efficiency in HCT-116 cells using immunoblotting and qRT-PCR.

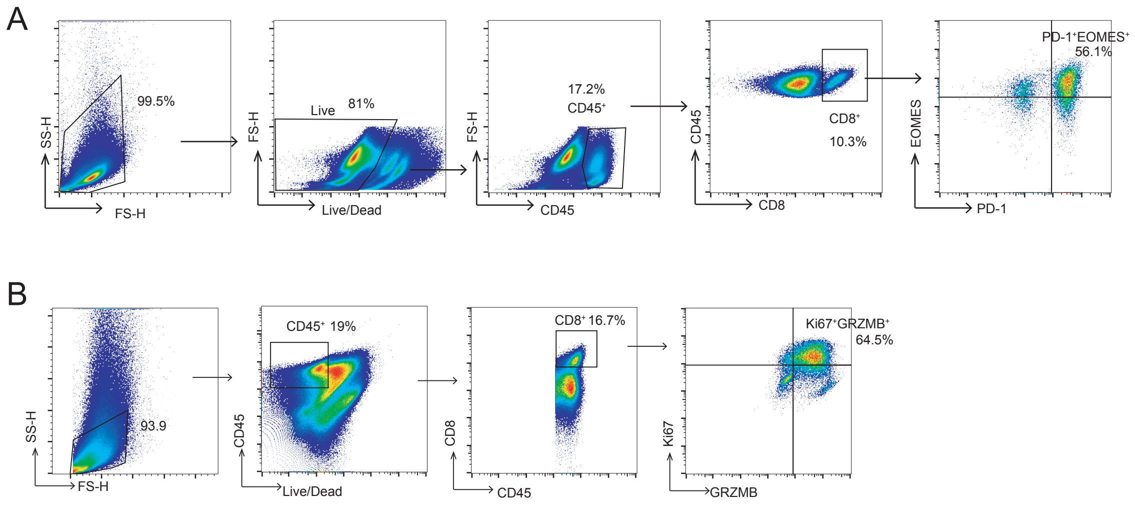


**Supplemental Figure 4.** Representative plots showing the gating strategy for the data of Figure 2I-2M (A) and Figure 3L (B).

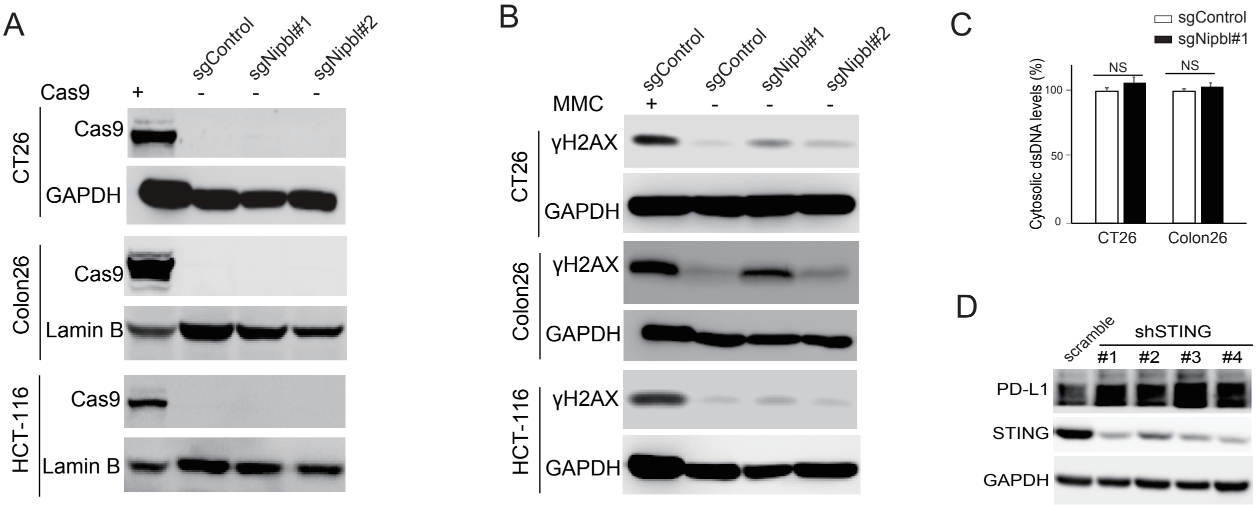


**Supplemental Figure 5.** The DNA damage is dispensable for Nipbl loss-induced tumour immunity. (A-B) The Cas9 and γH2AX were weakly or not expressed in sgNipbl subclones compared with sgControl cells. Exgenously expressed Cas9 and mitomycin C-induced γH2AX were used as positive controls.(C) The dsDNA content had no significant change in the cytosol of sgNipbl cells compared to that of sgControl. (D) shSTING did not affect PD-L1 expression in sgNIPBL#2 HCT-116 cells.

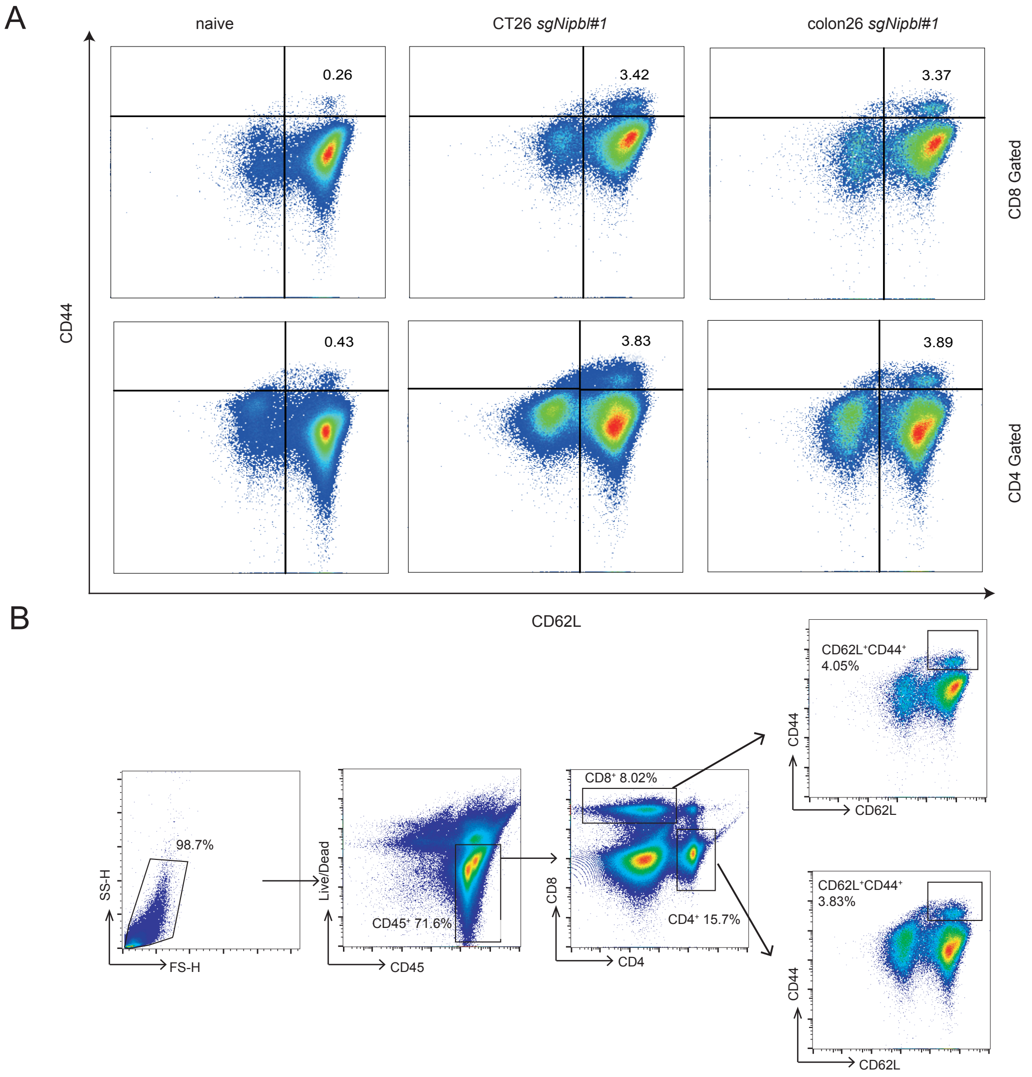


**Supplemental Figure 6.** Blockade of PD-1 elicited a potent anti-tumor memory response in host mice bearing sgNipbl tumors. (A) The representative plots of central memory CD8$^{+}$ and CD4$^{+}$ T cells showing CD62L$^{+}$ and CD44$^{+}$ in the spleens of naïve mice, sgNipbl #1 CT26 tumor-regressed mice and sgNipbl #1 colon26 tumor-regressed mice after anti-PD-1 therapy. (B) Representative plots showing the gating strategy for the data shown in (A).

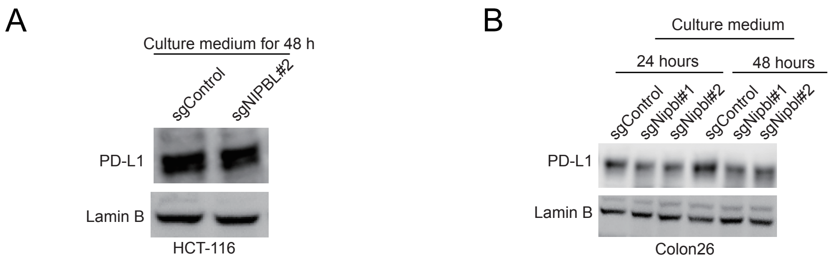


**Supplemental Figure 7.** The PD-L1 protein expression has little or no change upon sgNipbl cell culture medium treatment. (A) The PD-L1 protein expression levels of sgControl HCT-116 cells had no significant change after incubated with sgNipbl HCT-116 cell medium for 48 hours. (B) The PD-L1 protein expression levels of sgControl colon26 cells had no significant change after incubated with sgNipbl colon26 cell medium for indicated times. Lamin B were used as a loading control.

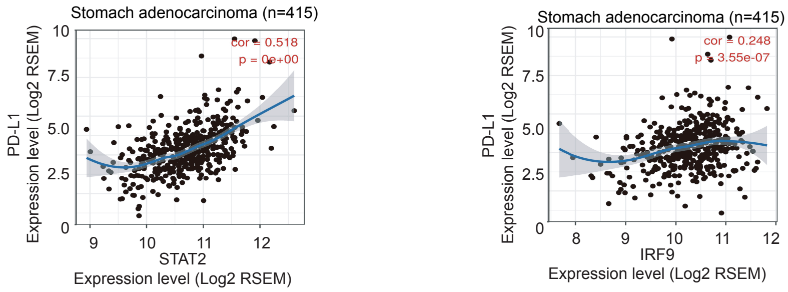


**Supplemental Figure 8.** The correlation of STAT2 and IRF9 mRNA levels with PD-L1 mRNA levels in TCGA stomach adenocarcinomas. The spearman's correlation and estimated significance were calculated by TIMER (Tumor Immune Estimation Resource) and adjusted for tumor purity.

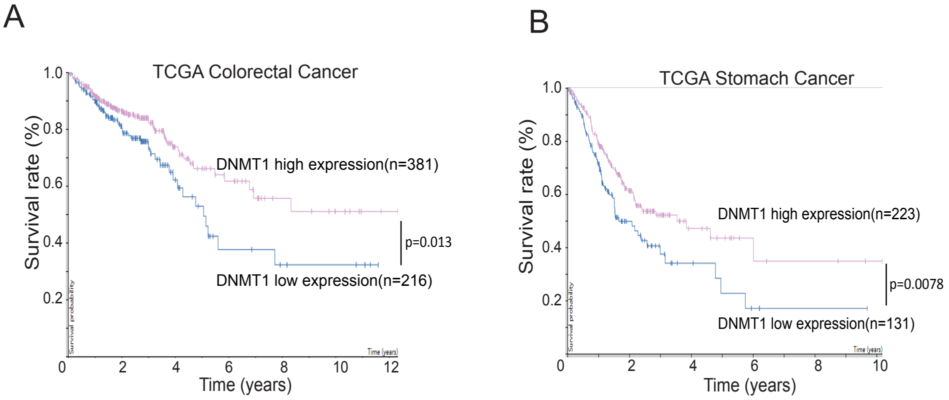


**Supplemental Figure 9.** Low DNMT1 mRNA levels are significantly correlated with the poor survival of TCGA stomach and colorectal cancer patients. The survival significance was assessed by a Log-rank Mantel-Cox test. Colorectal tumor samples, n=597; stomach tumor samples, n=354.

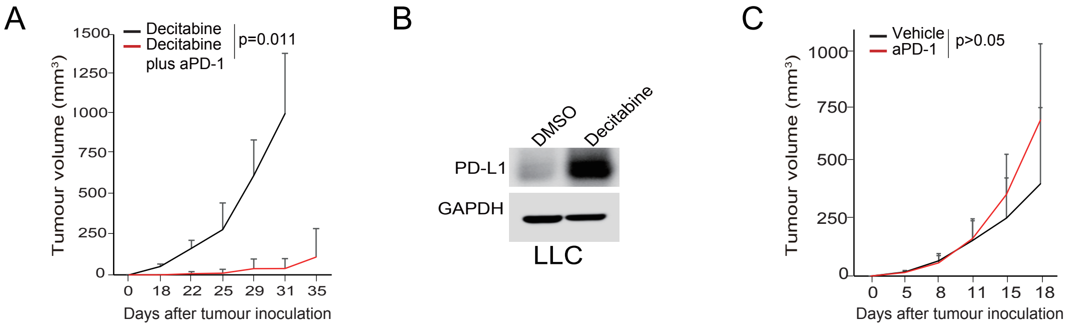


**Supplemental Figure 10.** PD-L1 is critical for the replase of DNMT1-inhibted lewis lung cancer cells in vivo. (A) Anti-PD1 monoclonal antibody potently inhibited the tumour growth in a decitabine-pretreated Lewis lung cancer syngeneic mouse model (decitabine-pretreated group: n=3; decitabine-pretreated plus anti-PD-1 group: n=3). Lewis lung cancer cells were pretreated with 0.5 μM decitabine for 3 days before inoculating into the flanks of syngeneic mice. (B) PD-L1 expression was determined after treated with 0.5 μM decitabine for 24 hours. (C) Anti-PD1 monoclonal antibody had no inhibitory effect on tumour growth in untreated Lewis lung cancer syngeneic mouse model (Vehicle: n=5, anti-PD-1: n=6).

a

| polytract site (wild-type) | Nucleotide change | Predicted amino acid change | Cancer patients incidence | Cell line incidence |
|---|---|---|---|---|
| AAAAAAAA | c.1808delA | K603Sfs*11 | 37 | 16 |
| AAAAAAA | c.8326delA | I2776Lfs*3 | 20 | 3 |
| TTTTTT | c.4269insT | V1424Cfs*6 | 8 | 0 |
| TTTTT | c.4269delT | F1424Lfs*15 | 5 | 1 |
| AAAAAAA | c.1513delA | R505Efs*35 | 5 | 0 |
| GGGGGG | c.2483insG | R949Qfs*17 | 3 | 0 |
| AAAAAAA | c.2252delA | N751Mfs*43 | 3 | 0 |
| AAAAAA | c.1697delA | K566Sfs*48 | 3 | 0 |
| CCCC | c.453insC | S152Lfs*21 | 2 | 0 |
| TTTTTT | c.6873delT | H2292Tfs*11 | 1 | 1 |
| AAAAA | c.7755delA | T2584Qfs*27 | 1 | 0 |
| AAAAAAA | c.2252insA | N751Kfs*2 | 1 | 0 |
| AAAA | c.5599insA | R1867Kfs*12 | 1 | 0 |
| AAAA | c.5599delA | R1867Efs*2 | 1 | 0 |
| AAAAAA | c.1697insA | P567Afs*2 | 1 | 0 |
| AAAAAA | c.7044delA | K2348Nfs*8 | 1 | 0 |
| AAAA | c.1163insA | N388Kfs*2 | 1 | 0 |
| TTTT | c.6659insT | L2220Ffs*3 | 1 | 0 |
| TTTT | c.4484insT | L1495Yfs*94 | 1 | 0 |
| GGGG | c.2506delG | D386Ifs*11 | 1 | 0 |
| AAAA | c.3747insA | A1250Sfs*7 | 1 | 0 |
| GGGGGG | c.2843delG | G948Afs*6 | 1 | 0 |
| AAAAAA | c.7044insA | Y2349Ifs*41 | 1 | 0 |
| GGGGG | c.1547delG | G948Afs*24 | 1 | 0 |
| AAAAAAA | c.4561delA | I1521Lfs*68 | 0 | 1 |
| AAAAAA | c.2328delA | K776Nfs*18 | 0 | 1 |
| AAAAAA | c.6706delA | N2236Tfs*29 | 0 | 1 |
| AAAAAAAA | c.1807delAA | K603Vfs*2 | 0 | 1 |
| AAAAA | c.5066insG | E1689Gfs*10 | 0 | 1 |

b

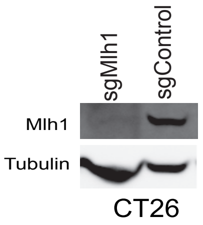


c

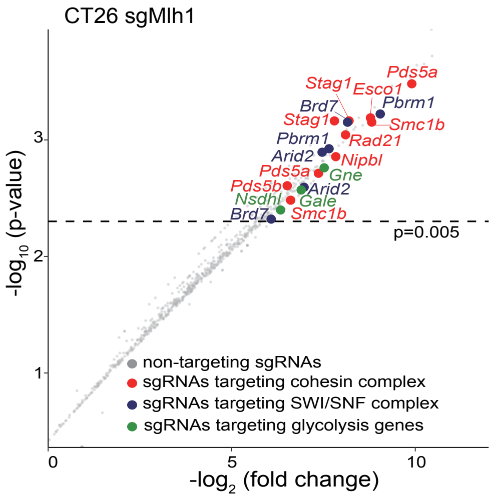


**Supplemental Figure 11.** sgRNAs targeting cohesin subunits and regulators is selectively enriched in the anti-PD-1 group of sgMlh1 CT26 tumor cells compared to that of vehicle group. (A) NIPBL are frameshift mutated at 29 polytract sites of 101 TCGA human tumor samples and 26 tumor cell lines. (B) The knockout efficiency of Mlh1 in CT26 were determined by immunoblotting. (C) sgRNA candidates that targets cohesin subunits and regulators (red), SWI/SNF complex (blue) and glycolysis (green) were presented. Dash indicates a p value of 0.005. Non-targeting sgRNAs (grey) was used as a negative control.

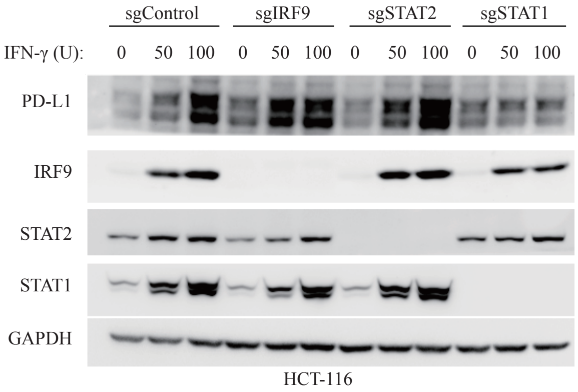


**Supplemental Figure 12.** IFN-γ induces PD-L1 expressin independent of STAT2 and IRF9. HCT-116 cells were treated with indicated concentrations of IFN-γ for 24 hours in the presence of sgControl, sgIRF9, sgSTAT2 and sgSTAT1, and the expression levels of indicated proteins were determined by immunoblottting.GAPDH was used as a loading control.

**Antibodies.** Primary antibodies against PD-L1 (#13684), STAT1 (#9172), p-STAT1 (Tyr701, #7649), p-STAT2 (Tyr690, #4441), IRF1 (#8478), p-IRF3 (#37829), RIG-I (#3743), MDA5 (#5321), MAVS (#3993), CDK4 (#12790), CDK6 (#13331), Cas9 (#14697), STING (#13647), NF-κB (#12540), p-c-Jun (Ser73, #3270), p-p38 (Thr180/Tyr182, #4511), p-MEK1/2 (Ser217/221, #9120), MLH1 (#3515) and GAPDH (#5174) were purchased from Cell Signaling Technology. Primary antibodies against NIPBL (sc-374625), MYC (sc-40), STAT2 (sc-1668), IRF3 (sc-33641), IRF7 (sc-74472), IRF9 (sc-365893), DNMT1 (sc-271729), vinculin (sc-25336) and lamin B (sc-6216) were purchased from Santa Cruz Biotechnology. Anti-dsRNA antibody (#MABE1134, clone rJ2) and γH2AX (#05-636) were purchased from Millipore Inc. Tubulin antibody (ab135209) was purchased from Abcam. Peroxidase-conjugated affiniPure goat anti-rabbit and goat anti-mouse second antibodies, Alexa Flour 488-conjungated goat anti-mouse/rabbit antibody and Rhodamine Red-conjugated goat anti-mouse/rabbit antibody were purchased from Jackson ImmunoResearch.

Flow cytometric antibodies against CD45 (Brilliant Violet 510™, #103138), CD8a (PerCP/Cy5.5, #100734), PD-L1 (PE, #123408), PD-1 (PE/Cy7, #135216), Ki-67 (FITC, #652409), Granzyme B (PE, # 372208), CD62L (Alexa Fluor® 647, #103026), CD44 (Alexa Fluor® 700, #103026), CD4 (FITC, #100510), CD25 (PE/Cy7, #102015), FOXP3 (Alexa Flour® 647, #126407), F4/80 (Alexa Flour® 647, #123122), CD11b (PerCP/Cy5.5, #101228), Ly6G/Ly6C (Gr-1) (Apc/Cy7, #108424), CD3 (PE, #100206) and NK-1.1 (Alexa Flour® 647, #108720) were purchased from Biolegend. Antibody against EOMES (eFluor 660, #50-4875-82) was purchased from ThermoFisher Scientific.

**Plasmids.** Mouse GeCKO library V2 (#1000000053), pRSV-Rev (#12253), pMDLg/pRRE (#12251), pMD2.G (#12259), LentiCRISPR v2 (#52961), pLX304 (#25890), pLKO.1-puro (#8453) and pLenti CMV Puro DEST (#17452) plasmids were purchased from Addgene. The cDNAs of STAT1a, STAT2, IRF1, IRF3, IRF7, IRF9 and MAVS were cloned from HEK-293T cells with indicated primers, and were further constructed into pLX-304 or pLenti CMV Puro DEST vectors with sequencing verification. Primers used in gene cloning were listed as follows:

STAT1_F: 5’-CGCGGATCCATGTCTCAGTGGTACGAACTT-3’;

STAT1_R: 5’-ATAAGAATGCGGCCGCCTATACTGTGTTCATCATACTGT-3’;

STAT2_F: 5’-CGCGGATCCGCTCATACTAGGGACGGGAAG-3’;

STAT2_R: 5’-tttGCTCTTCGtataTTTGACCGTGAAGCTGATGATGTGCA-3’;

IRF1_F: 5’-CGCGGATCCATGCCCATCACTCGGATGC-3’;

IRF1_R: 5’-ATAAGAATGCGGCCGCCTACGGTGCACAGGGAAT-3’;

IRF3_F: 5’- GCCGGTACCATGGGAACCCCAAAGCCAC-3’;

IRF3_R: 5’- GCCCTCGAGTTATTGGTTGAGGTGGTGGGGAAC-3’;

IRF7_F: 5’-CGCGGATCCATGGCCTTGGCTCCTGA-3’;

IRF7_R: 5’-ATAAGAATGCGGCCGCCTAGGCGGGCTGCTCCAGCTCCAT-3’;

IRF9_F: 5’-CGCGGATCCATGGCATCAGGCAGGGCA-3’;

IRF9_R: 5’-ATAAGAATGCGGCCGCCTACACCAGGGACAGAATG-3’;

MAVS_F: 5’- tccggtaccgGCCACCatgccgtttgctgaagacaagacct-3’;

MAVS_R: 5’- ctaCTTGTCATCGTCGTCCTTGTAGTCgtgcagacgccgccggtacagcacca ccaggagtgtgactaccagcacccct-3’.

**Flow cytometry analysis.** Tumour tissue was cut into small pieces with a scissor, followed by treatment with collagenase IV (Roche) and DNase I (Roche) in RPMI 1640 for 20 min at 37°C. Cell suspension was filtered through a 40 μm filter in PBS. Live/dead cell discrimination was performed using LIVE/DEAD ™ Fixable Violet Dead Cell Stain Kit (ThermoFisher Scientific, Cat#L34955). Cell surface staining was done for 30 min at 4°C. Intracellular staining was performed with Foxp3/Transcription Factor Staining Buffer Set (ThermoFisher Scientific, Cat#00-5523-00). All flow cytometric analysis was conducted using Gallios (Beckman) and analysed using FlowJo software.

**Quantitative PCR.** Total RNA was extracted with Trizol reagent and cDNA was synthesized using PrimerScript RT reagent Kit. Quantitative PCR was performed on Applied Biosystem 7500 Real-time PCR system using NovoStart SYBR supermix. *GAPDH* was used as an internal control to normalize input of cDNA. Primers for real-time PCR were listed as follows:

Human_GAPDH_F: 5’-AAGGTGAAGGTCGGAGTCAA-3’;

Human_GAPDH_R: 5’-AATGAAGGGGTCATTGATGG-3’;

Human_CD274_F: 5’-CAAAGAATTTTGGTTGTGGA-3’;

Human_CD274_R: 5’-AGCTTCTCCTCTCTCTTGGA-3’;

Human_IRF9_F: 5’-GCCCTACAAGGTGTATCAGTTG-3’;

Human_IRF9_R: 5’-TGCTGTCGCTTTGATGGTACT-3’;

Human_IRF1_F: 5’-CTGTGCGAGTGTACCGGATG-3’;

Human_IRF1_R: 5’-ATCCCCACATGACTTCCTCTT-3’;

Human-SMC1A_F: 5’- CCTGAGACCTTCTTGCCTCTTG-3’;

Human-SMC1A_R: 5’- GAGGTGGCTCATAGCGAATCAC-3’;

Human_STAT2_F: 5’-GAGCCAGCAACATGAGATTGA-3’;

Human_STAT2_R: 5’-GCCTGGATCTTATATCGGAAGCA-3’;

Human_STAT1_F: 5’-TTACTCCAGGCCAAAGGAAG-3’;

Human_STAT1_R: 5’-TTCAGCTGTGATGGCGATAG-3’;

Human_DNMT1_F: 5’- CCTAGCCCCAGGATTACAAGG-3’;

Human_DNMT1_R: 5’- ACTCATCCGATTTGGCTCTTTC-3’;

Human_HERVE_F: 5’- GGTGTCACTACTCAATACAC-3’;

Human_HERVE_R: 5’- GCAGCCTAGGTCTCTGG-3’;

Human_HERVF_F: 5’- CCTCCAGTCACAACAACTC-3’;

Human_HERVF_R: 5’- TATTGAAGAAGGCGGCTGG-3’;

Human_HERVK_F: 5’- ATTGGCAACACCGTATTCTGCT-3’;

Human_HERVK_R: 5’- CAGTCAAAATATGGACGGATGGT-3’;

Human_RIG-I_F: 5’- CCAGCATTACTAGTCAGAAGGAA-3’;

Human_RIG-I_R: 5’- CACAGTGCAATCTTGTCATCC-3’;

Human_MDA5_F: 5’- GAGCAACTTCTTTCAACCACAG-3’;

Human_MDA5_R: 5’- CACTTCCTTCTGCCAAACTTG-3’;

Human_IFNα_F: 5’- AGAAGGCTCCAGCCATCTCTGT-3’;

Human_IFNα_R: 5’- TGCTGGTAGAGTTCGGTGCAGA-3’;

Human_IFNß_F: 5’-ATGACCAACAAGTGTCTCCTCC-3’;

Human_IFNß_R: 5’- GGAATCCAAGCAAGTTGTAGCTC-3’;

Human_IFNγ_F: 5’- TCGGTAACTGACTTGAATGTCCA-3’;

Human_IFNγ_R: 5'- TCGCTTCCCTGTTTTAGCTGC-3';

Human_IL28_F: 5'- AGGGCCAAAGATGCCTTAGA-3';

Human_IL28_R: 5'- TCCAGAACCTTCAGCGTCAG-3';

Human_IL29_F: 5'- GGACGCCTTGGAAGAGTCAC-3';

Human_IL29_R: 5'- AGCTGGGAGAGGATGTGGT-3';

Human_IRF3_F: 5'- AGAGGCTCGTGATGGTCAAG -3';

Human_IRF3_R: 5'- AGGTCCACAGTATTCTCCAGG -3';

Human_IRF7_F: 5'- GCTGGACGTGACCATCATGTA -3';

Human_IRF7_R: 5'- GGGCCGTATAGGAACGTGC-3';

Mouse_Cd274_F: 5'-GCTCCAAAGGACTTGTACGTG-3';

Mouse_Cd274_R: 5'-TGATCTGAAGGGCAGCATTTC-3';

Mouse_Gapdh_F: 5'-AGGTCGGTGTGAACGGATTTG-3';

Mouse_Gapdh_R: 5'-TGTAGACCATGTAGTTGAGGTCA-3'.

**CRISPR-Cas9 mediated knockout.** $2\times10^5$ cells were transfected with 1 μg lenti-CRISPR-v2 vector with indicated sgRNA using lipofectamine 3000 (Invitrogen) following the manufacturer's guidelines. After puromycin selection, transfected cells were plated into 96-well-plate with a density of less than 1 cell/well. The knockout effects of indicated genes in each subclone were validated by immunoblotting and Sanger sequencing. sgRNAs used in this study were listed as follows:

Mouse_Nipbl sgRNA#1, 5'-AACCAAATTGTCATCCCTAC-3';

Mouse_ Nipbl sgRNA#2, 5'-GTGTTGTGTAACTTGAAGGA-3';

Mouse_ Nipbl sgRNA#3, 5'-CGATATACCCGTCTTGTTGC-3';

Mouse_Stag2 sgRNA#1, 5'-AACCAAATTGTCATCCCTAC-3';

Mouse_Stag2 sgRNA#2, 5'-GTGTTGTGTAACTTGAAGGA-3';

Mouse-Mlh1 sgRNA, 5'-CTAATTCAGATCCAAGACAA-3';

Non-targeting sgRNA, 5'- GGGTCTTCGAGAAGACCT-3'

Human_NIPBL sgRNA, 5'-TCCCCATTACTACTCTTGCG-3';

Human_IRF9 sgRNA#1, 5'-GGGCCCCCCACCGAAGTTCC-3';

Human_IRF9 sgRNA#2, 5'- CAGCAACTGATACACCTTGT-3';

Human_STAT2 sgRNA#1, 5'-CTCTGTGCAACCGTACACGA-3';

Human_STAT2 sgRNA#2, 5’-TGAGATTGAATCCCGGATCC-3’;

Human_STAT1 sgRNA#1, 5’-GTGGAGCGGTCCCAGAACGG-3’;

Human_STAT1 sgRNA#2, 5’-GAGGTCATGAAAACGGATGG-3’;

Human_IRF1 sgRNA#1, 5’-GAACTCCCTGCCAGATATCG-3’;

Human_IRF1 sgRNA#2, 5’-TTAATTCCAACCAAATCCCG-3’.

**shRNA mediated RNA interference.** Cells were infected with lentiviruses containing indicated shRNAs supplemented with 8 μg/ml polybrene. After replacement with fresh medium, cells were selected by puromycin in order to remove uninfected cells. The knockdown efficiency was determined by immunoblotting or qPCR with indicated primers. shRNAs used in this study was listed as follows;

SMC1A_shRNA#1, 5’-CCAACATTGATGAGATCTATA-3’;

SMC1A_shRNA#2, 5’-CGGCGTATTGATGAAATCAAT-3’;

STAT1_shRNA#1, 5’-CAAGCGTAATCTTCAGGATAA-3’;

STAT1_shRNA#2, 5’-ACAGCTGTTACTCAAGAAGAT-3’;

STAT1_shRNA#3, 5’-ACCTTGCAGAACAGAGAACAC-3’;

STAT1_shRNA#4, 5’-GAACAGAAATACACCTACGAA-3’;

IRF1_shRNA#1, 5’-CAGATTAATTCCAACCAAATC-3’;

IRF1_shRNA#2, 5’-GCGTGTCTTCACAGATCTGAA-3’;

IRF3_shRNA#1, 5’- GATCTGATTACCTTCACGGAA-3’;

IRF3_shRNA#2, 5’- GGGAAGAGTGGGAGTTCGAGG-3’;

IRF7_shRNA#1, 5’- CGCCTAGAACCCAGTCTAATG-3’;

IRF7_shRNA#2, 5’- GCTGGACGTGACCATCATGTA-3’;

IRF7_shRNA#3, 5’- GACATCGAGTGCTTCCTTATG-3’;

MAVS_shRNA#1, 5’- ATGTGGATGTTGTAGAGATTC-3’;

MAVS_shRNA#2, 5’- CCAGAGGAGAATGAGTATAAG-3’;

MAVS_shRNA#3, 5’- GGAGAGAATTCAGAGCAAGCC-3’;

STING_shRNA#1, 5’-GGTCATATTACATCGGATATC-3’;

STING_shRNA#2, 5’- GCAGAGCTATTTCCTTCCACA-3’;

STING_shRNA#3, 5’- GGGCACCTGTGTCCTGGAGTAC-3’;

STING_shRNA#4, 5’- GCCCGGATTCGAACTTACAAT-3’.

**RNA sequencing.** The cells were lysed in Trizol reagent, and total RNAs were exacted for RNA sequencing accomplished by Novogen Company using Illumina HiSeq X Ten platform. Sequenced reads were aligned to Ensemble database with STAR. Gene expression values were normalized to obtain RPKM values using RSEM.

**Immunoprecipitation.** Cell lysates were incubated with 1 μg indicated primary antibody at 4°C overnight, followed by addition of 10 μl PurePtoteome$^{TM}$ protein A/G Mix Magnetic Beads magnetic beads (Millipore) for 2 h at 4°C. After washed with lysis buffer for 5 times, the precipitates were incubated with elution buffer and heated at 70°C on a shaker for 10 min. The lysates were then mixed with 4× Laemmle buffer, and were subjected to immunoblotting. ChromPure Mouse IgG (015-000-003, Jackson ImmunoResearch) was used as an internal control.

**Mice genotyping and spontaneous tumor models.** All the animal experiments were performed according to the procedures approved by the Institutional Animal Care and Use Committee, Shanghai Institute of Nutrition and Health. The Nipbl$^{fl/fl}$ mouse (B6; 129S-Nipbl$^{tm1(flox)Smoc}$, #NM-CKO-00067) was purchased from Shanghai Model Organisms. The Apc$^{Min}$ mice was a kind gift from Prof. Jun Qin. Villin-Cre mouse (T002376, B6/JNju-Tg(Pvillin-cre)I/Nju) was purchased from NRCMM (Nanjing, China). To obtain Nipbl conditional knockout mice in gastrointestinal epithelial cells, Nipbl$^{fl/fl}$ mouse was first crossed with wild-type C57/B6 mice to remove neo gene and then crossed with Villin-Cre mouse. To get Apc$^{Min}$; Nipbl$^{fl/fl}$, the Villin-Cre+; Nipbl$^{fl/fl}$ mouse was crossed with Apc$^{Min}$ mouse. Mice were marked and genotyped about 10 days after born. Briefly, tail tip was boiled for 20 min in 200 μL of 0.05M NaOH, then neutralized with 20 μL Tris-EDTA buffer (Tris 1M, pH 8, EDTA 10 mM), serving as PCR template. The genotype of Nipbl$^{fl/fl}$ mouse was validated by the size of PCR products using following primers: F, 5'-ATGTTGTGAAATTAAGTTCCTAAG-3', R, 5'-GTCAAAATACTATATGCTTTCTTT-3'. For the loxP site in intron 4 of the Nipbl$^{fl/fl}$ allele, the PCR products were 300 bp for the floxed allele and 330 bp for the wild-type allele. Detection of the Cre yielded a 350-bp fragment.

Mice were separated into three groups with indicated numbers after genotyped, termed as Nipbl$^{fl/fl}$ (Nipbl HO), Villin-Cre+; Nipbl$^{fl/+}$ mice (Nipbl HE;Cre) and Villin-Cre+;

$Nipbl^{fl/fl}$ (Nipbl HO;Cre). When crossed with $Apc^{Min}$, mice were divided into two groups, termed as $Apc^{Min}$ (Apc) and $Apc^{Min}$; Villin-Cre+; $Nipbl^{fl/fl}$ (Nipbl HO;Cre/Apc) with indicated numbers after genotyped.

**Immunofluorescence staining.** Appropriate density of cells were seed on glass coverslips for 36 hours. Cells on glass coverslip were gently washed by 1×PBS for 3 times and were then fixed by 4% Paraformaldehyde plus 0.2% Triton X100 for 15 min, following for 3 times again. After blocked with 3% BSA for 40 min, cells were stained with indicated primary antibody for 2 hours and were then subjected to 1×PBS washing for 5 times. After incubated with secondary antibody for 40 min, cells were stained with DAPI for 10 min. The glass coverslips were mounted, and images were captured by OLYMPUS FV1200. For quantification of cytosolic dsRNA, at least 100 cells from each group were scored.

**Cytosolic dsRNA extraction and quantification.** For cytosolic dsRNA extraction, cells were lyzed by 1×cell lysis buffer containing 20 mM HEPES PH7.9, 10 mM KCL, 0.5 mM spermidine and 0.1% Triton X100. After centrifugation, total cytosolic dsRNAs in the supernatants were extracted using TRIzol reagent and incubated with RNaseA in high salt concentration for 10 mins at room temperature. After extracted by TRIzol again, the concentration of cytosolic dsRNA were determined by Nanodrop 2000c or Qbit.

**ChIP-PCR.** After crosslinked by formaldehyde, cells were subjected to chromatin-immunoprecipitation following the guidelines of the SimpleChIP Plus Enzymatic Chromatin IP Kit (CST, Magnetic Beads, #9005). Normal IgG was used as negative control. ChIP-PCR Primers for DNMT1 promoter were listed as follows: F, 5’-ACCTCCAGCTTTCCCATTTT-3’, R, 5’- TTGAGCTGCCTGATCACATC-3’. ChIP-PCR Primers for PD-L1 promoter were listed as follows: F, 5’-AAAGCCATATGGGTCTGCTG-3’, R, 5’-CAACATCTGAACGCACCTTG-3’.

**Luciferase Reporter Assays.** The 1 kb PD-L1 promoter fragment was cloned into pGL3-basic-luciferase vector using the primers as follows: F, 5’-GCTTCTTTTGAGTGGGCAGA-3’, R, 5’-AATTGTGGAAACCCAACAGG-3’. The

60-bp STAT2 and IRF9 binding motifs were removed by a second round of PCR. The DNMT1 promoter fragment was cloned into pGL3-basic luciferase vector using the primers as follows: F, 5’-ACTGCTAAACCTATATC-3’, R, 5’- GTCAGGAGTTCTGG ACCAGC-3’. The STAT2 promoter fragment was cloned into pGL3-basic luciferase vector using the primers as follows: F, 5’- AACGTAAGAACGTAAATGTTTGGGTG TTGAAGTCACAGGGTTTGGTTTCGA-3’, R, 5’- CTGATTAGGGTTGCAGTCCC CGCGCCCTCCAATGGCTCTGGTCGCGACTTCCCGTCCCTAGTATGAGCT-3’. The IRF9 promoter fragment was cloned into pGL3-basic luciferase vector using the primers as follows: F, 5’- AAGCTGACAGAAGAGGTACCCTTGGGACA-3’, R, 5’- TGACCTTTGCCTACAAGGGTCTGGGGAGCTGGCCCTTGTCAGTCTCACTC-3’. Cells were transfected with 500 ng pGL3-PD-L1-luciferase and 50 ng pRL-TK vectors using calcium phosphate transfection method. The luciferase expression levels were measured using a Dual Luciferase® Reporter Assay System (Promega), and the relative luciferase ratio was calculated using Renilla luciferase as an internal control.
